# Broad remodeling of the pulmonary immune landscape occurs during IAPA with specific functional deficits of neutrophil subsets

**DOI:** 10.64898/2026.07.10.737810

**Authors:** Nima Naghshtabrizi, Dequan Lou, Yonne Karoline Tenório de Menezes, Ravineel B. Singh, Shuxia Wang, Caden Ngeow, Fan Li, Radha Gopal, Kong Chen, Keven Robinson

**Affiliations:** Division of Pulmonary, Allergy, Critical Care and Sleep Medicine, Department of Medicine, University of Pittsburgh, Pittsburgh, Pennsylvania, USA; University of Pittsburgh School of Medicine, Department of Medicine, Pittsburgh, Pennsylvania, USA; Department of Pediatrics, UPMC Children’s Hospital of Pittsburgh, Pittsburgh, Pennsylvania, United States

## Abstract

**Background:** Influenza-associated pulmonary aspergillosis (IAPA) is a severe complication of influenza infection associated with prolonged intensive care unit stay and increased mortality. Although impaired antifungal immunity has been implicated in IAPA pathogenesis, the cell type-specific immune mechanisms driving susceptibility remain incompletely understood. We aimed to characterize the pulmonary immune landscape during IAPA using single-cell transcriptomics and functional neutrophil assays.

**Methods:** Male C57BL/6 mice were assigned to naïve control, influenza A/PR/8/34 (H1N1) infection, *A. fumigatus* (ATCC 42202) infection, or IAPA groups. Lung CD45^+^ immune cells underwent single-cell RNA sequencing with downstream clustering and CellChat ligand-receptor interaction analysis. Differential gene expression analyses were performed across myeloid, lymphoid, and neutrophil populations. Functional neutrophil responses were evaluated using flow cytometry, myeloperoxidase activity assays, and FLARE (fluorescent Aspergillus reporter) conidia to assess fungal conidia uptake and killing. Cross-species validation was performed using gene set enrichment analysis compared with published human IAPA transcriptomic datasets.

**Results:** IAPA broadly remodeled the pulmonary immune landscape across myeloid, lymphoid, and neutrophil compartments. Myeloid cells showed coordinated suppression of fungal pattern recognition receptors, lysosomal biogenesis programs, and inflammatory signaling. Lymphoid populations exhibited transcriptional signatures of T cell exhaustion and Th17 suppression. Within the neutrophil compartment, we identified two transcriptionally and functionally distinct populations, conventional and inflammatory neutrophils, with divergent antifungal effector capacities. Inflammatory neutrophils showed selective killing defects, while both subsets exhibited impaired phagocytic uptake during IAPA. Murine transcriptomic findings demonstrated strong concordance with immune dysfunction signatures identified in human IAPA.

**Conclusion:** IAPA susceptibility arises from coordinated transcriptional dysfunction spanning innate and adaptive immune compartments. Distinct neutrophil subset dysfunction, impaired fungal recognition pathways, and T-cell exhaustion signatures collectively contribute to defective fungal clearance, providing mechanistic insight into IAPA susceptibility and potential therapeutic targets.

## Introduction

Influenza infection represents a substantial public health concern, seasonally affecting millions worldwide and causing between 290,000 and 650,000 respiratory-related mortalities annually^1^. Influenza predisposes patients to the development of secondary pneumonia, including both bacterial and fungal superinfections. Recent studies suggest that up to 19% of critically ill patients hospitalized with influenza will develop influenza-associated pulmonary aspergillosis (IAPA). The diagnosis of IAPA can be challenging, and IAPA typically results in prolonged intensive care unit (ICU) stays and increased mortality^2^. Effective host defense against both influenza viruses and *Aspergillus fumigatus* (*A. fumigatus*) necessitates a coordinated response from both the innate and adaptive immune systems. The innate immune system serves as a rapid initial line of defense in the lungs, where resident alveolar macrophages, epithelial cells, and neutrophils identify inhaled pathogens through pattern-recognition receptors. These cells respond by engulfing the invaders and releasing inflammatory cytokines and reactive oxygen species (ROS) ^3^. Dendritic cells are activated to present antigens to T lymphocytes, thereby establishing a connection between the innate immune response and adaptive immunity. In turn, the adaptive immune system provides targeted and sustained protection, with CD4⁺ helper T cells enhancing the microbicidal activity of macrophages and neutrophils, and CD4⁺ T helper 17 cells promoting neutrophil recruitment and antifungal defense through IL-17 and IL-22 signaling ^3–5^. To gain a deeper understanding of the immunopathology that drives IAPA, we utilized single-cell transcriptomics to facilitate an in-depth analysis of cell type-specific responses, allowing for the identification of dysregulated pathways and immune dysfunction that occurs during IAPA. Neutrophils are critical effectors of antifungal immunity, and emerging evidence has revealed that neutrophils comprise transcriptionally and functionally distinct subpopulations whose composition shifts dynamically in response to infection^6^. We leveraged our single-cell transcriptomic data to examine neutrophil subset effector function during IAPA.

## Methods

### Animal Models

Male C57BL/6 wild-type mice aged 6–8 weeks were obtained from Taconic Farms. All animals were maintained under specific pathogen-free conditions with protocols approved by the University of Pittsburgh Institutional Animal Care and Use Committee.

### Pathogens and IAPA model

C57BL/6 mice were challenged with influenza A/PR/8/34 (10^2^ pfu) or PBS control for 6 days prior to *Aspergillus fumigatus* (ATCC strain 42202, 2.5×10^7^ resting conidia) or PBS control. Five experimental groups were established as illustrated in Figure 1A: PBS naïve control, influenza Day 6, influenza Day 8, *A. fumigatus* alone, and IAPA coinfection. All experimental groups were harvested at day 8, except for the influenza Day 6 group which was harvested on Day 6 post-influenza infection. Influenza A/PR/8/34 (H1N1) was kindly provided by Dr. Radha Gopal (UPMC Children’s Hospital of Pittsburgh). Mice were inoculated with 100 plaque-forming units (PFU) in 50 µL sterile PBS via oropharyngeal aspiration.

**Figure 1.**
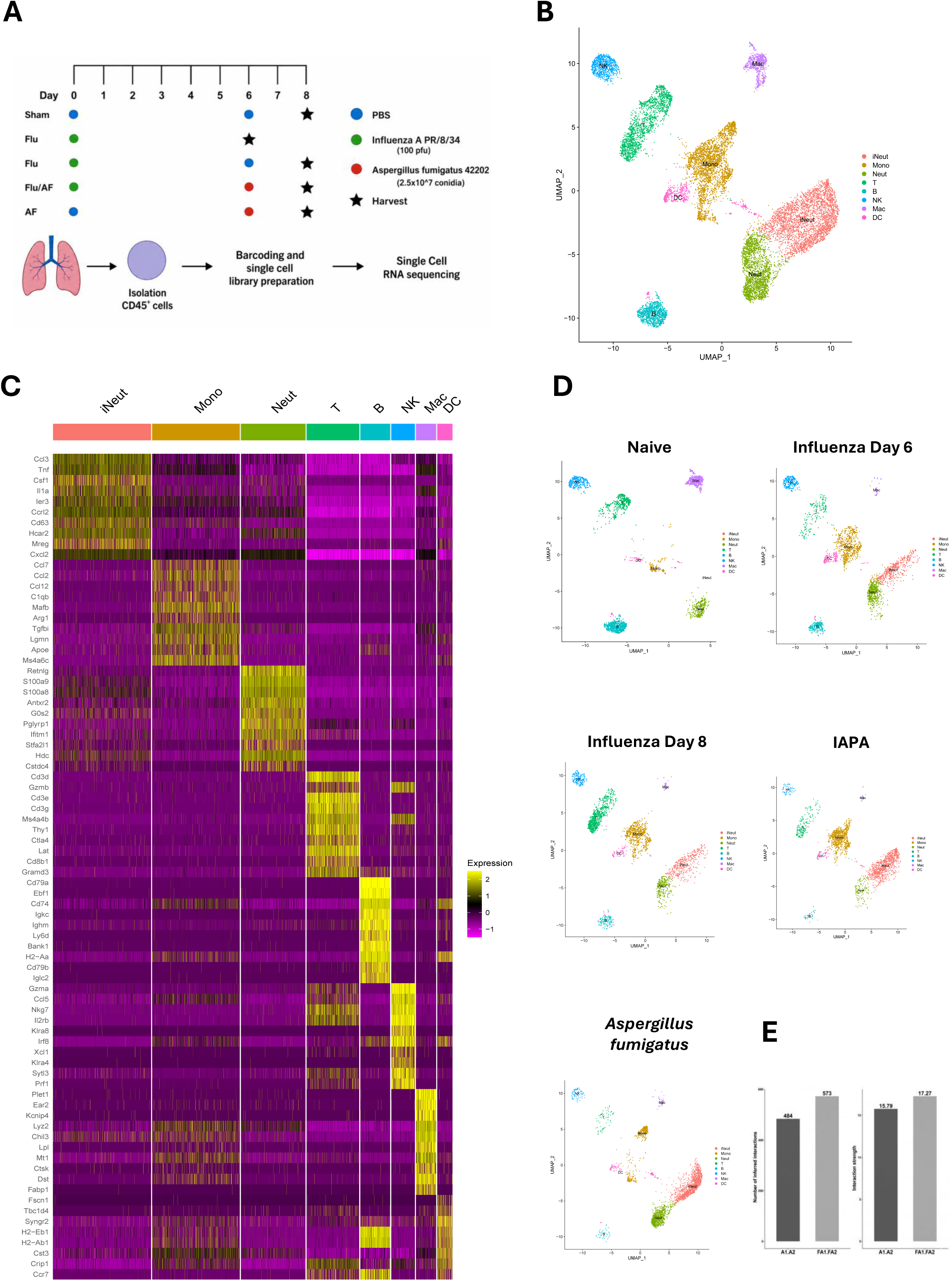
Cellular composition of lung immune cells during IAPA. **(A)** Schematic of experimental design and workflow. Two mice per experimental group were used for scRNA-seq analyses. **(B)** UMAP plot of all 10 integrated samples **(C)** Heatmap of marker genes corresponding to each identified cluster **(D)** UMAP plot of all samples, divided by treatment groups **(E)** Bar charts showing the total number of inferred ligand–receptor interactions (left) and aggregate interaction strength (right) comparing *Aspergillus fumigatus* infection (A1.A2) and IAPA (FA1.FA2).

*A. fumigatus* ATCC 42202 was propagated on potato dextrose agar at 37°C for 5–7 days. Conidia were harvested by washing culture plates with sterile PBS containing 0.1% Tween 20 and enumerated using a hemocytometer. Mice were challenged with 2.5 × 10^7^ conidia in 50 µL sterile PBS via oropharyngeal aspiration on day 6.

The <u>fl</u>uorescent *<u>A</u>spergillus* <u>re</u>porter (FLARE) strain Af293-dsRed was generously provided by Dr. Tobias Hohl (Memorial Sloan Kettering Cancer Center) and prepared as previously described^7^. Briefly, 1 × 10^8^ conidia were incubated with 0.5 mg/mL Biotin-XX SSE (ThermoFisher) in 50 mM carbonate buffer (pH 8.3) at 4°C for 2 hours under continuous rotation, washed with 0.1 M Tris-HCl, and conjugated with 0.02 mg/mL streptavidin–Alexa Fluor 647 (AF-647; Life Technologies) for 30 minutes at room temperature protected from light. Labeled conidia were resuspended in PBS containing 0.025% Tween 20 and stored at 4°C until use.

### Single-Cell RNA Sequencing

CD45⁺ immune cells were isolated from mouse lungs, 2 mice per experimental group, using CD45 MicroBeads (Miltenyi Biotec) and processed for single-cell RNA sequencing (scRNA-seq) (Figure 1A). Reads were aligned to the mouse reference genome (mm10/GRCm38) and quantified with Cell Ranger (v7.0; 10x Genomics), and the filtered feature–barcode matrices were imported into R with Read10X. Samples were multiplexed using cell-hashing antibodies; hashtag oligo (HTO) counts were added as a separate assay and normalized by centered log-ratio (CLR) transformation. Cells were demultiplexed with HTODemux (positive quantile = 0.99), and negative droplets and cross-sample doublets were removed, retaining only singlets. All downstream processing was performed in Seurat (v5.2.0). A Seurat object was created retaining genes detected in ≥3 cells and cells with ≥200 detected genes, and low-quality cells were further excluded by retaining only cells with 200–6,000 detected genes and <5% mitochondrial reads. Counts were log-normalized, highly variable features were identified, and the data were scaled prior to principal component analysis. Cells from all samples were analyzed together as a single integrated dataset; the first 15 principal components were used to construct a shared nearest-neighbor graph and cells were clustered with the Louvain algorithm (resolution = 0.2) and visualized by UMAP. Cluster identities were assigned from canonical lineage markers as inflammatory neutrophils, conventional neutrophils, monocytes, macrophages, dendritic cells, T cells, B cells, and NK cells; a small mixed T–monocyte cluster was excluded from comparative analyses. Cluster markers were identified with FindAllMarkers (Wilcoxon rank-sum test). Myeloid, lymphoid, and neutrophil compartments were subset and re-clustered to resolve substructure. Between-condition differential gene expression within each cell type was performed with FindMarkers using the MAST test. For general pathway analyses, gene set enrichment analysis (GSEA) was performed with fgsea against the MSigDB mouse hallmark gene sets (msigdbr; Mus musculus; category H), with genes ranked by average log₂ fold-change.

### Cell Line Authentication/Validation Statement

No cell lines were used to perform the experiments within this manuscript.

### Gene Set Enrichment Analysis and Cross-Species Concordance

To assess transcriptomic concordance between our murine IAPA model and human IAPA, GSEA was performed on cell-type-specific differential gene expression comparing IAPA versus singular influenza infection, mirroring the primary biological contrast reported in Feys et al ^8^. GSEA was conducted using the fgsea package in R with pathway gene sets curated to include pathways reported as downregulated in human IAPA bronchoalveolar lavage samples by Feys et al., at either the pathway enrichment or gene level of evidence. Enrichment scores were calculated across all eight immune cell populations identified by scRNA-seq. Statistical significance was determined using Benjamini-Hochberg adjustment, with a threshold of p < 0.25 applied as recommended for GSEA analyses. Normalized enrichment scores (NES) were visualized as a heatmap across cell types and pathways.

### Cell–Cell Communication Analysis (CellChat)

Intercellular communication was inferred with CellChat (v2.1.2). For each condition, a CellChat object was created from the log-normalized expression matrix using the annotated cell-type labels as identities. Ligand–receptor pairs were drawn from the mouse CellChat database (CellChatDB.mouse, “Secreted Signaling” category). Over-expressed signaling genes and interactions were identified, and expression values were projected onto the mouse protein–protein interaction network. Communication probabilities were computed with computeCommunProb, and interactions supported by fewer than 10 cells in either the sending or receiving population were removed (filterCommunication, min.cells = 10). Pathway-level probabilities were aggregated (computeCommunProbPathway, aggregateNet) and network centrality was computed to define signaling roles. To compare A. fumigatus and IAPA conditions, the two objects were merged (mergeCellChat) and contrasted in the total number and overall strength of inferred interactions (compareInteractions), the signaling roles of each cell type, the relative information flow of individual pathways (rankNet), and differential ligand–receptor signaling (netVisual_bubble).

### Flow Cytometry

Lungs were harvested into 1 mL sterile PBS and processed immediately. Tissue was digested in DMEM media containing Collagenase IV (1 mg/mL) and DNase I (0.1 mg/mL) under continuous rotation at 37°C for 1 hour. Following digestion, samples were mechanically dissociated and passed through a 70-µm cell strainer. Red blood cells were lysed using ACK lysis buffer nd the remaining cell pellet was resuspended in 1 mL PBS. Cells were stained with a surface antibody cocktail containing: CD45 (BUV496, clone 30-F11, BD OptiBuild), Ly6G (BV510, clone 1A8, BioLegend), and CD11b (BUV395, clone M1/70, BD Horizon) for 30 minutes at 4°C protected from light. Following surface staining, cells were permeabilized and stained intracellularly with an antibody cocktail containing: IL-1α (StarBright Violet 440, clone ALF-161, BioLegend, #503202) conjugated in-house using the TrailBlazer Tag Kit (Bio-Rad, #12020038) and TrailBlazer StarBright Violet 440 Label Kit (Bio-Rad, #12020042) according to the manufacturer’s instructions, and TNF-α (BV786, clone MP6-XT22, Invitrogen, #417-7321-82) for 20 minutes at room temperature protected from light. Samples were washed and acquired immediately on a Cytek Aurora spectral flow cytometer and analyzed using FlowJo software (BD Biosciences). Gating strategies for all populations are provided in Figure S1-Supplemental File 1. To determine absolute cell counts, the number of events acquired for each population by flow cytometry was normalized to the total cell count obtained by automated cell counting prior to staining, accounting for the fraction of cells loaded per well relative to the total cell suspension.

## Results

### Single-cell transcriptomic profiling reveals broad remodeling of the pulmonary immune landscape during IAPA

To characterize the immune response across distinct models of pulmonary infection, we performed scRNA-seq on CD45⁺ immune cells isolated from the lungs of mice infected with singular influenza infection, singular *A. fumigatus* infection, IAPA, and naïve controls (Figure 1A). Unsupervised UMAP clustering of the integrated dataset revealed distinct immune cell populations that were annotated based on canonical lineage markers (Figure 1B–C, Figure S2A-Supplemental File1). The abundance and distribution of specific immune cell populations varied across different infection models, demonstrating that cellular composition of immune cells in the lung is dynamically remodeled across all infection conditions, with the most pronounced changes observed during IAPA (Figure 1D and Table S1-Supplemental File 1). For each immune cell cluster, heatmaps showing all differentially expressed genes across treatment conditions are provided (Figure S3, A-C, Supplemental File 1). CellChat analysis of ligand–receptor interactions across all immune cell populations revealed a greater number of inferred interactions and higher aggregate interaction strength during IAPA compared to singular A. *fumigatus* infection (573 vs 484 interactions; 17.27 vs 15.79 aggregate strength), suggesting that prior influenza infection broadly remodels intercellular communication networks in the lung at the time of secondary fungal challenge (Figure 1E). Comprehensive ligand-receptor interaction networks for all remaining immune cell populations are provided in Supplemental File 2.

### IAPA drives coordinated suppression of myeloid pattern recognition and antifungal effector programs

Among myeloid populations (Figure 2A), macrophage, monocyte, and dendritic cell populations were identified based on marker gene expression (Figure 2B, Figure S2B-Supplemental File 1). Unsupervised UMAP clustering of the integrated dataset revealed distinct myeloid cell populations whose composition varied across experimental groups (Figure 2C). Macrophage abundance was highest in naïve mice and progressively decreased across infection conditions, with the most pronounced reduction observed during IAPA, suggesting that prior influenza infection broadly alters myeloid cell composition in the lung (Table S1). Naïve macrophages were characterized by upregulation of *Chil3*, *Lyz2*, and *Plet1* (Figure 2B). Monocyte abundance was highest during IAPA relative to all other infections, and monocytes were characterized by upregulation of *Ccl2*, *Ccl7*, and *Ccl12*, key chemokines involved in monocyte recruitment and infiltration during inflammation^9,10^ (Figure 2B and Table S1). Dendritic cells, which showed comparable numbers across infection models, exhibited high expression of *Cts3* and *Ccr7*, a chemokine receptor that directs dendritic cell migration from peripheral tissues to lymph nodes to initiate adaptive immune responses ^11^ (Figure 2B and Table S1). We initially used an unbiased approach to identify the top differentially expressed genes in myeloid cell clusters (Figure 2D). Next, we compared gene expression between singular *A. fumigatus* infection and IAPA to help determine the genes most changed during *A. fumigatus* infection with and without preceding influenza infection (Figure 2E). In macrophages, *Tfeb* and *Tyk2*, key regulators of lysosomal biogenesis and IL-10–mediated immunosuppression ^12–14^, respectively, alongside *Cxcl3*, a neutrophil chemoattractant^15^, were downregulated during IAPA compared to singular *A. fumigatus* infection, while the anti-inflammatory and antibody-dependent phagocytic mediators *Il10*, *Tnfaip6*, and *Fcgr1* (CD64) were upregulated ^16,17^ (Figure 2E).

**Figure 2.**
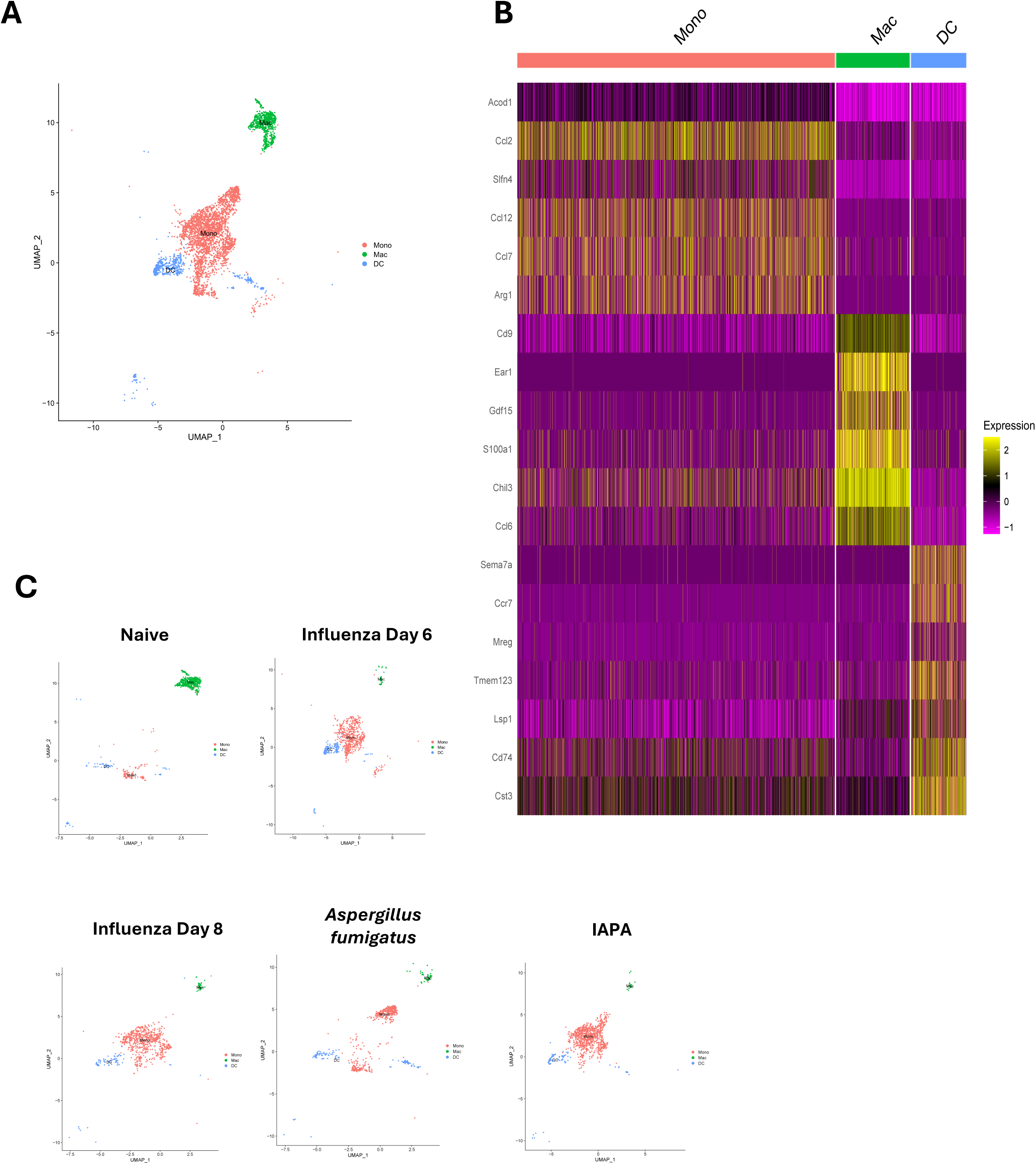
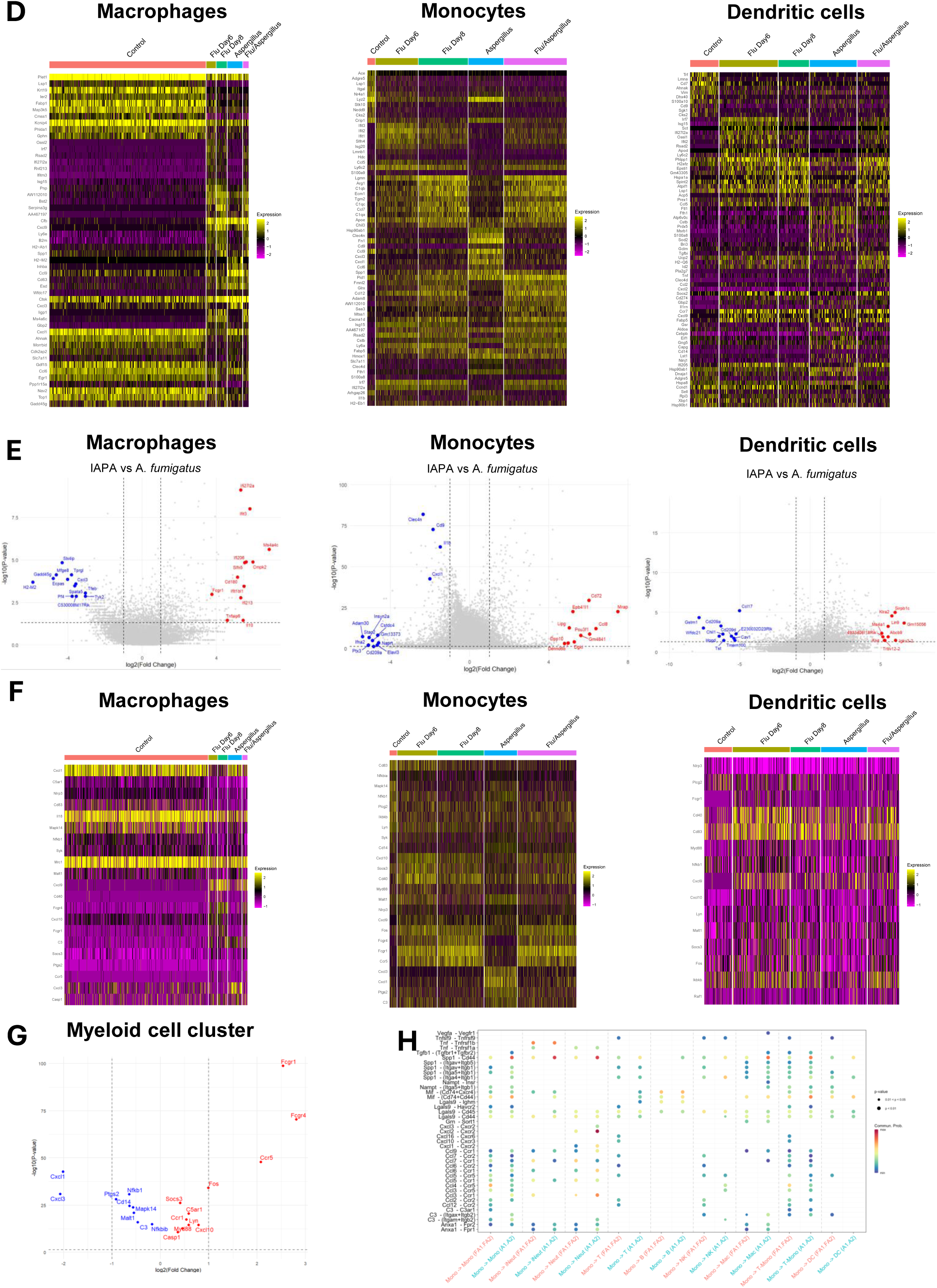
IAPA drives coordinated suppression of myeloid pattern recognition and antifungal effector programs. Influenza Day 6 mice were harvested at day 6; all remaining groups were harvested at day 8. **(A)** UMAP plot of all integrated samples **(B)** Heatmap of biologically relevant differentially expressed genes in each myeloid cluster **(C)** UMAP plot of all samples, divided by treatment groups **(D)** Differential gene expression in myeloid clusters (unbiased approach - 10 most upregulated genes and 10 most downregulated genes for each treatment group) **(E)** Volcano plot of the top differentially expressed genes in myeloid cell populations (Comparing IAPA to *Aspergillus fumigatus* alone) **(F)** Differential gene expression of anti-fungal genes in myeloid clusters **(G)** Volcano plot of the top differentially expressed antifungal genes in myeloid cell population (Comparing IAPA to *Aspergillus fumigatus* alone) **(H)** Ligand–receptor interactions originating from monocytes. Dot plot showing inferred ligand–receptor interactions with monocytes as the source cell population, comparing *Aspergillus fumigatus* infection (A1.A2) and IAPA (FA1.FA2) across all target cell types. Dot size reflects p-value significance, and color reflects communication probability.

*Clec4n* (Dectin-2), a C-type lectin receptor that binds to galactomannan in the cell wall of *A. fumigatus*, was downregulated in monocytes during IAPA compared singular *A. fumigatus* infection^18^ (Figure 2E). Similarly, Pentraxin 3 (*Ptx3*), a key soluble recognition pattern receptor that recognizes *A. fumigatus,* was downregulated during IAPA compared with singular *A. fumigatus* infection (Figure 2E). Monocyte-derived genes involved in neutrophil chemotaxis (*Cxcl1*), the inflammasome pathway (*Il1b*), and leukocyte adhesion and migration (*Cd9*) were also downregulated during IAPA relative to singular *A. fumigatus* ^19,20^. In dendritic cells, *Cd209a* and *Cd209d*, which bind *A. fumigatus* conidia via high-mannose structures including galactomannan, were downregulated during IAPA compared to singular *A. fumigatus* infection^21^. *Ccl17*, which reduces recruitment of activated dendritic cells and macrophages ^22^, was downregulated during IAPA compared with singular *A. fumigatus* infection (Figure 2E). Finally, we focused on transcription of antifungal genes in myeloid cells (Figure 2F-G) and the pattern recognition receptor genes *Cd14* and *C3* were downregulated in IAPA compared with singular *A. fumigatus* infection (Figure 2G). Additionally, *Cxcl1* and *Cxcl3* expression were significantly reduced in myeloid cells during IAPA compared with singular *A. fumigatus* infection (Figure 2G), consistent with previously reported reductions in total CXCL1 and CXCL2 levels in the lungs at 6 and 24 hours after *A. fumigatus* challenge during IAPA, which were associated with decreased neutrophil infiltration ^23^. Both *Fcgr1* and *Fcgr4*, receptors that bind immunoglobulin antibodies, and *Ccr5*, which binds cytokines, were significantly upregulated in myeloid cells during IAPA compared to singular *A. fumigatus* infection (Figure 2G). Ligand–receptor interactions originating from monocytes revealed distinct intercellular communication patterns between *A. fumigatus* and IAPA conditions across all immune cell populations (Figure 2H). Together, these data demonstrate coordinated transcriptional suppression of antifungal effector programs across macrophage, monocyte, and dendritic cell populations during IAPA.

### Conventional and inflammatory neutrophils exhibit selective transcriptional and functional defects during IAPA

Neutrophils are the primary effectors of antifungal immunity against *A. fumigatus*, and prior influenza infection has been shown to impair both pulmonary neutrophil recruitment and function during secondary *A. fumigatus* challenge ^23,24^. To determine whether transcriptionally distinct neutrophil subpopulations contribute to these defects, we identified two transcriptionally distinct neutrophil populations: conventional neutrophils and inflammatory neutrophils (Figure 3A). We defined inflammatory neutrophils as IL-1α and TNFα positive based on the scRNA-seq because these cells showed enriched expression of canonical pro-inflammatory cytokines and associated inflammatory gene programs. IL-1α and TNFα were chosen as defining markers because they were among the most differentially expressed and biologically relevant inflammatory mediators within the overall neutrophil cluster, providing a robust and interpretable way to distinguish these calls from other neutrophil states present in the dataset. Compared to conventional neutrophils, inflammatory neutrophils demonstrated high expression of *Tnf*, *Il1a*, *Cd63*, and *Ccl3*. Conventional neutrophils demonstrated high expression of *S100a8*, *S100a9*, and *Mmp8* (Figure 3B, Figure S2C-Supplemental File 1). When comparing across infectious groups, inflammatory neutrophils were essentially absent in the PBS naïve control group. During influenza infection, the neutrophil cluster transitioned from both conventional neutrophils and inflammatory neutrophils at day 6 post-infection to higher numbers of inflammatory neutrophils at day 8 post-infection. Both conventional and inflammatory neutrophil populations were lower in abundance during IAPA compared to *A. fumigatus* alone, consistent with previously reported influenza-mediated suppression of pulmonary neutrophil recruitment in some IAPA models ^23,25^ (Figure 3C and Table S1-Supplemental File 1). Although the number of inflammatory neutrophils was reduced during IAPA compared to *A. fumigatus* infection, the larger and more significant difference was observed in the conventional neutrophil group, which had severely reduced abundance during IAPA compared to *A. fumigatus* infection. Using an unbiased approach to identify the top differentially expressed genes in neutrophil clusters, we observed that the antimicrobial peptide gene, *Lcn2* was upregulated in inflammatory neutrophils during IAPA compared to singular *A. fumigatus* infection, while conventional neutrophils showed the opposite pattern, with *Lcn2* downregulated (Figure 3D–E). Interferon-signaling genes, *Ifit1*, *Ifit2*, *Ifit3*, *Rsad2*, *Isg15*, and *Isg20*, were upregulated in IAPA compared to *A. fumigatus* in both the conventional and inflammatory neutrophil populations (Figure 3D–E). *Il1f6* and *Il1f9*, which bind the IL-36 receptor and enhance Th1 and Th17 responses via antigen-presenting cells^26^, were downregulated in inflammatory neutrophils during IAPA compared to *A. fumigatus* (Figure 3E). Next, we focused on transcription of antifungal genes in neutrophil populations (Figure 3F) and observed that *Nfkb1*, *Syk*, and *Nlrp3*, which are activated by the hyphal form of *A. fumigatus* and drive inflammasome assembly and caspase-1–mediated IL-1β maturation^14^, were downregulated during IAPA compared to *A. fumigatus* infection, while *Nfkbia* was upregulated^27^ (Figure 3G). CellChat analysis further revealed distinct ligand–receptor interaction patterns originating from inflammatory neutrophils between *A. fumigatus* and IAPA conditions (Figure 3H). Together, these transcriptional and functional data demonstrate that IAPA is associated with the emergence of two neutrophil subpopulations with divergent antifungal effector capacities, characterized by impaired conidial uptake across both subsets and selective killing defects within the inflammatory neutrophil population.

**Figure 3.**
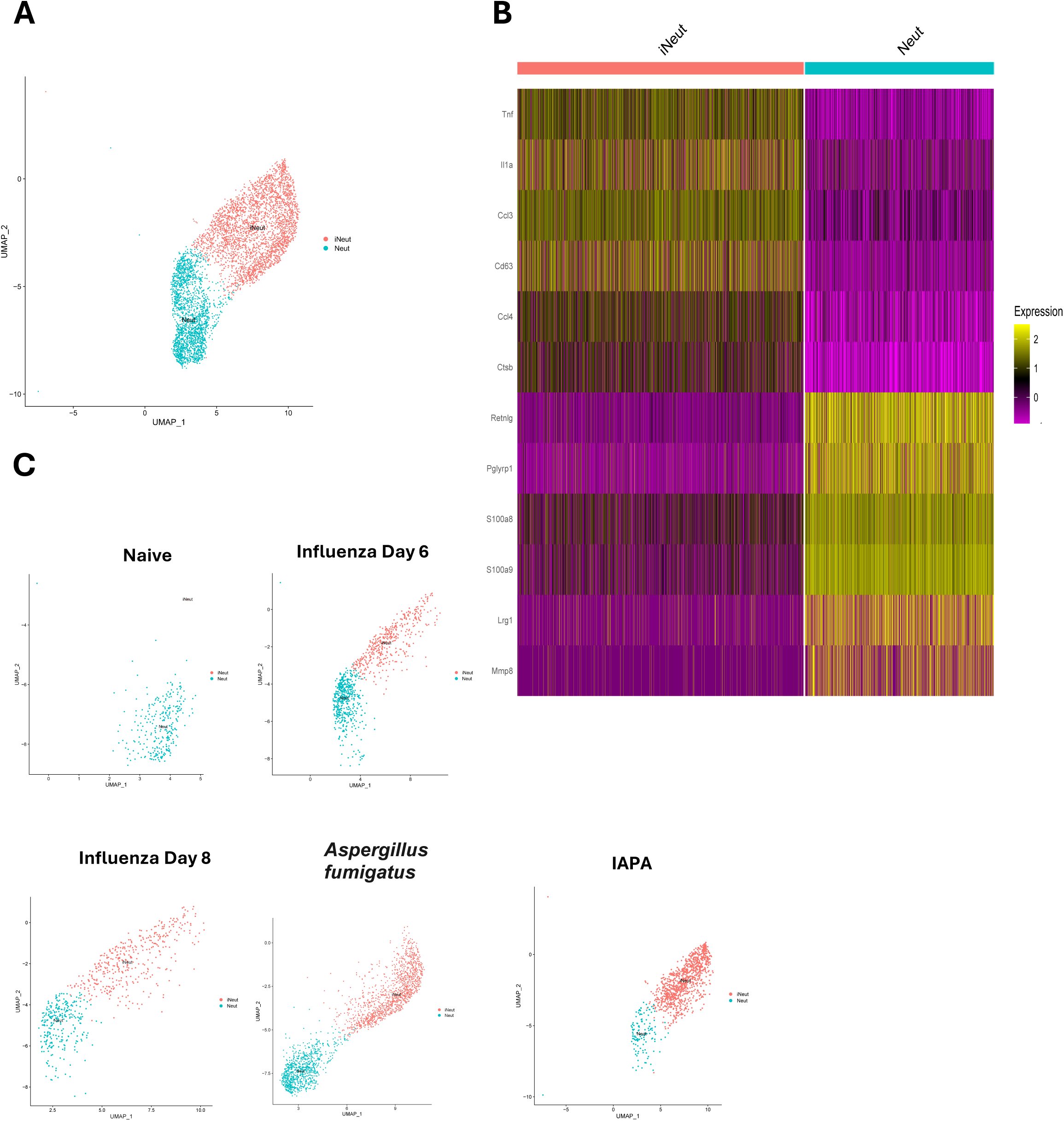
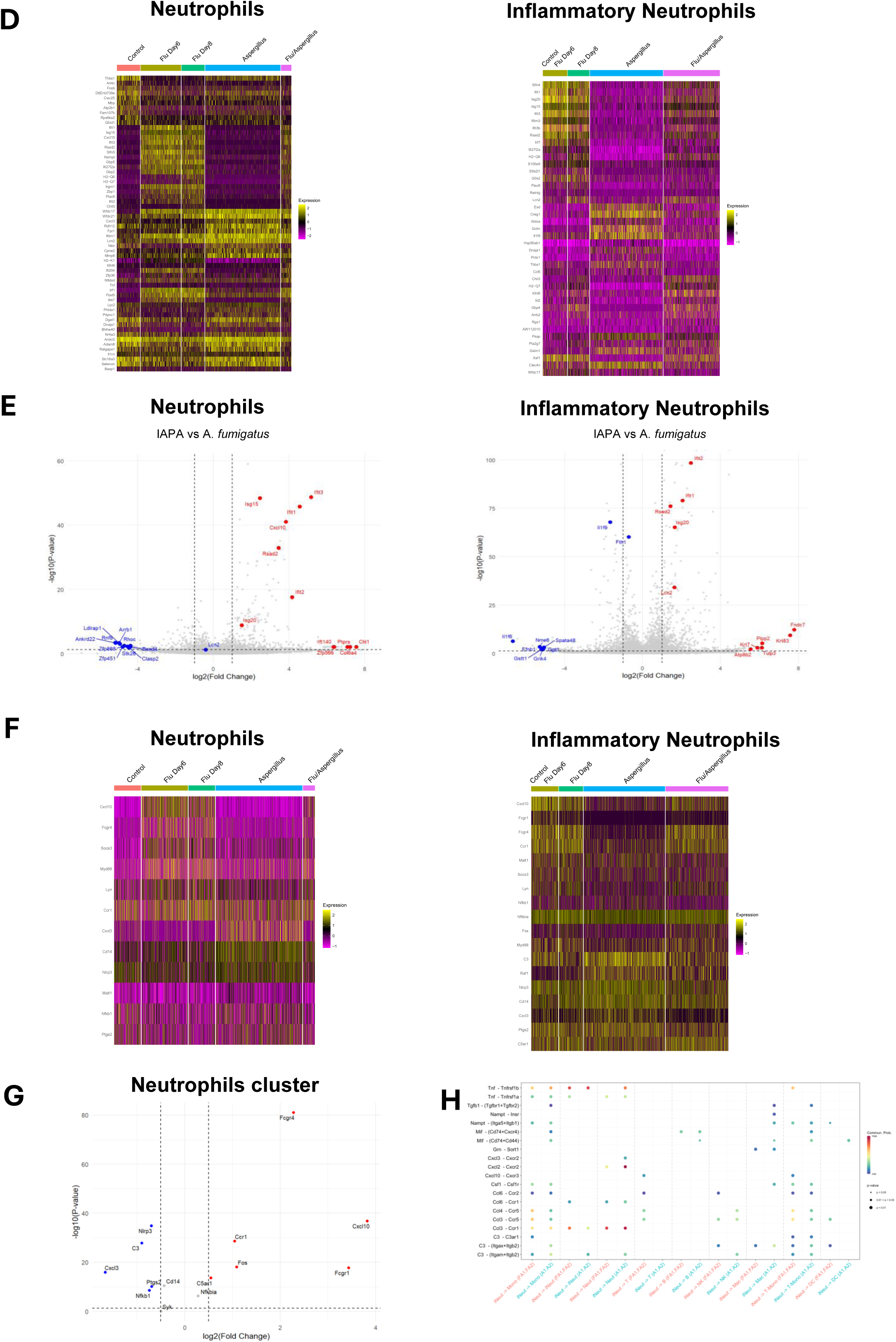
Conventional and inflammatory neutrophils exhibit selective transcriptional and functional defects during IAPA. Influenza Day 6 mice were harvested at day 6; all remaining groups were harvested at day 8. **(A)** UMAP plot of all integrated samples **(B)** Heatmap of biologically relevant differentially expressed genes in each neutrophil cluster **(C)** UMAP plot of all samples, divided by treatment groups **(D)** Differential gene expression in neutrophil clusters (unbiased approach - 10 most upregulated genes and 10 most downregulated genes for each treatment group) **(E)** Volcano plot of the top differentially expressed genes in neutrophil cell populations (Comparing IAPA to *Aspergillus fumigatus* alone) **(F)** Differential gene expression of anti-fungal genes in neutrophil clusters **(G)** Volcano plot of the top differentially expressed antifungal genes in neutrophils **(H)** Ligand–receptor interactions originating from Inflammatory Neutrophils (iNeut). Dot plot showing inferred ligand–receptor interactions with iNeut as the source cell population, comparing *Aspergillus fumigatus* infection (A1.A2) and IAPA (FA1.FA2) across all target cell types. Dot size reflects p-value significance, and color reflects communication probability.

### Coordinated suppression of cytokine-driven innate responses and neutrophil effector function impairs fungal clearance during IAPA

To characterize neutrophil effector functions during IAPA, we isolated neutrophils from the lungs of mice infected with influenza, *A. fumigatus,* and IAPA and measured myeloperoxidase (MPO) enzyme activity. MPO is stored in neutrophil granules and released when neutrophils encounter *A. fumigatus* to activate microbial killing. We observed decreased MPO activity during IAPA compared to singular influenza or *A. fumigatus* infection (Figure 4A). To validate transcriptional findings at the protein level and further characterize neutrophil effector functions during IAPA, we utilized FLARE conidia ^7^ in our model and observed increased fungal burden during IAPA compared to *A. fumigatus* infection, similar to prior studies (Figure 4B)^25,28^. Consistent with increased fungal burden, IAPA mice exhibited significantly greater weight loss compared to mice infected with *A. fumigatus* over the course of infection (Figure S4-Supplemental File 1). FLARE conidia express DsRed as an indicator of fungal viability and were coupled to the dye Alexa Fluor 647 (AF647), acting as a signal independent of viability. Live FLARE conidia emit AF647 and DsRed and can be visualized within neutrophils. FLARE conidia that have been taken up and killed by neutrophils emit AF647 but not DsRed, which is turned off in the phagolysosome (Figure S4-Supplemental File 1). We used flow cytometry to identify total neutrophils (LiveCD45^+^CD11b^+^Ly6G^+^), conventional neutrophils (LiveCD45^+^CD11b^+^Ly6G^+^IL-1α^lo^TNFα^lo^), and inflammatory neutrophils (LiveCD45^+^CD11b^+^Ly6G^+^IL-1α^high^TNFα^high^). We observed a reduction in total neutrophils during IAPA compared to *A. fumigatus* infection at 48 hours post-fungal challenge (Figure 4C), similar to what we and others have previously published ^23,25,29^. These neutrophils exhibited similar killing capacity between IAPA and *A. fumigatus* mice, but a significant reduction in phagocytic uptake during IAPA compared to *A. fumigatus* (Figure 4D-E). Next, we examined the neutrophil subsets identified as conventional neutrophils and inflammatory neutrophils using our scRNA-seq dataset and observed a reduction in the total numbers of conventional neutrophils during IAPA compared to singular *A. fumigatus* (Figure 4F). Although there was a trend towards a reduction in inflammatory neutrophils during IAPA compared to singular *A. fumigatus*, the difference was not significant. These findings are similar to the total numbers of cells in the clusters observed in the scRNA-seq dataset (Table S1). We observed a reduction in killing capacity in IAPA inflammatory neutrophils compared to *A. fumigatus* inflammatory neutrophils, but no differences in killing capacity in the conventional neutrophil groups (Figure 4G). Both conventional and inflammatory neutrophil subsets exhibited significantly reduced conidial uptake during IAPA relative to their counterparts in singular *A. fumigatus* infection (Figure 4H).

**Figure 4.**
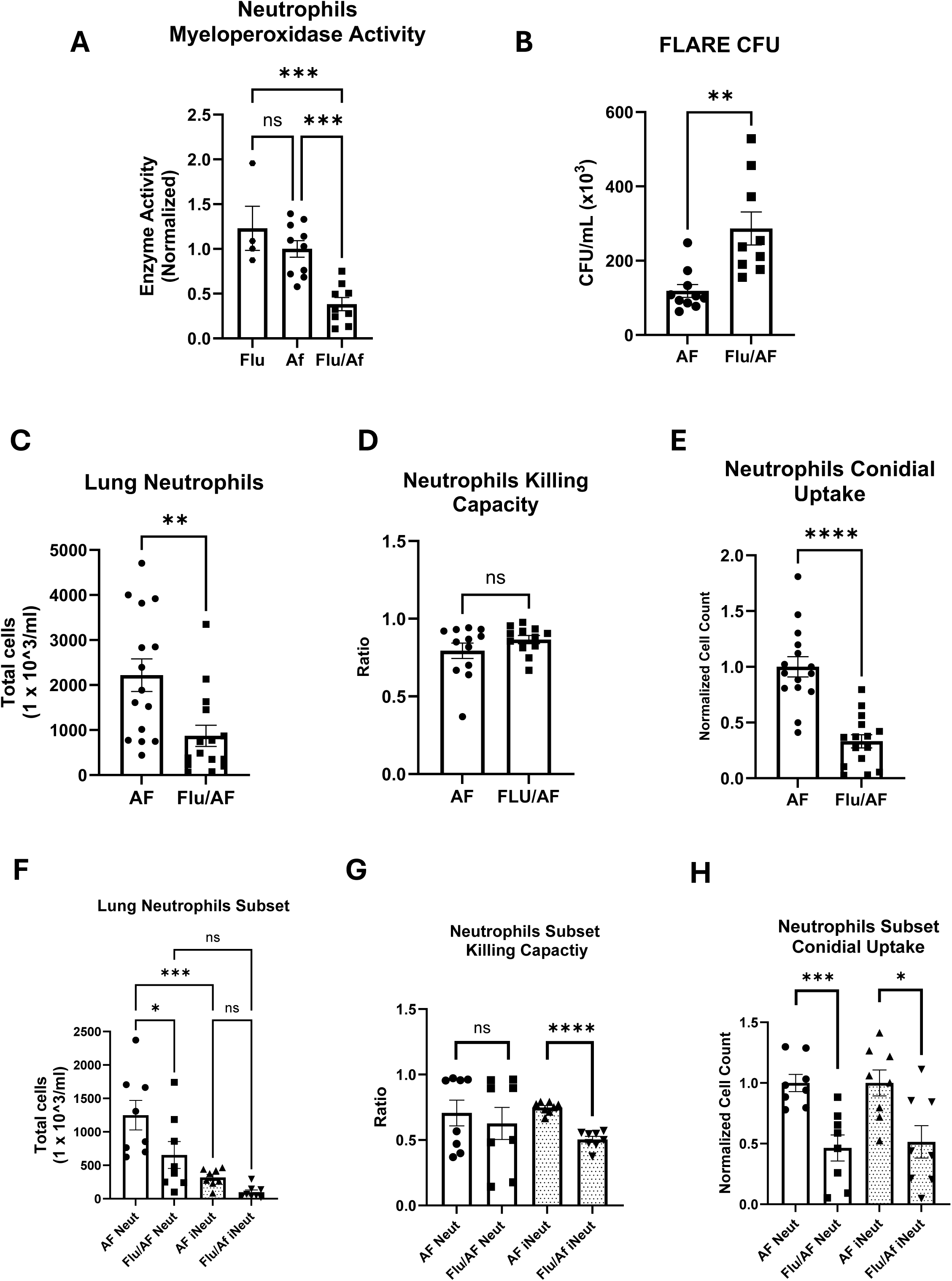
Defects in Neutrophil Antifungal Function During IAPA. Mice were infected with <u>fl</u>uorescent <u>A</u>spergillus <u>re</u>porter (FLARE) conidia in the presence or absence of prior influenza infection. Lungs were harvested at day 2 post-fungal challenge, single-cell suspensions were prepared, and neutrophil subsets were analyzed by flow cytometry. **(A)** Myeloperoxidase enzyme activity of neutrophils isolated from lung homogenate of Flu, AF, and Flu/AF groups, normalized to control. **(B)** Fungal burden (CFU/mL) of *Aspergillus fumigatus Af293* in lung homogenates of AF and Flu/AF groups at day 2 post-challenge. **(C)** Total lung neutrophil counts in AF and Flu/AF groups. **(D)** Conidial killing capacity of neutrophils in AF and Flu/AF groups, expressed as a ratio. **(E)** Total conidial uptake by neutrophils in AF and Flu/AF groups. **(F)** Total counts of conventional (Neut) and inflammatory (iNeut) neutrophil subsets in AF and Flu/AF groups. **(G)** Killing capacity of Neut and iNeut subsets in AF and Flu/AF groups. **(H)** Conidial uptake by Neut and iNeut subsets in AF and Flu/AF groups. Data were compiled from two independent experiments and are presented as means ± SEM. *p<0.05, **p<0.005, ***p<0.0005, ****p<0.0001 by one-way and two-way ANOVA with Tukey’s multiple comparisons test (ns, not significant).

### Prior influenza infection disrupts lymphoid antifungal immunity through T cell exhaustion

Among lymphoid populations (Figure 5A), T cell, B cell, and natural killer (NK) cell populations were identified based on marker gene expression (Figure 5B-Figure S2D-Supplemental File 1). Unsupervised UMAP clustering of the integrated dataset revealed lymphoid cell populations that varied between infection (Figure 5C). T lymphocytes, characterized by high expression of *Cd3d* and *Lat*, were increased at day 8 post-influenza infection but comparatively reduced in both singular *A. fumigatus* and IAPA, both of which groups were harvested at the same time point. T lymphocyte numbers increased significantly from day 6 post-influenza infection to day 8 post-influenza infection (Figure 5C, and Table S1). B lymphocytes, characterized by expression of *Ebf1*, *Cd79a,* and *Bank1*, were reduced in number in both IAPA and *A. fumigatus* infection compared to influenza infection (Figure 5C, and Table S1). NK cells, marked by high expression of *Nkg7*, *Klra8*, and *Prf1*, were reduced in number in both IAPA and *A. fumigatus* infection compared to influenza infection (Figure 5C and Table S1). We initially used an unbiased approach to identify the top differentially expressed genes in lymphoid cell clusters (Figure 5D). Next, we compared gene expression between singular *A. fumigatus* infection and IAPA to help determine the genes most changed during *A. fumigatus* infection with and without preceding influenza infection (Figure 5E). *Lgals3* (Galectin-3) was downregulated in NK cells across all infection models compared to control, consistent with reports of impaired neutrophil extravasation and higher fungal burden in galectin-3–deficient mice during pulmonary *A. fumigatus* infection (Figure 5D) ^30^. As described previously, Type 17 immune-related genes, *Il17a*, *Il17f*, *Il22*, and *Il23r*, were downregulated in IAPA compared to *A. fumigatus* infection ^23,31^. *Pdcd1* and *Lag3*, key regulators of T cell exhaustion and Treg suppressive function^32,33^, showed particularly elevated expression in both IAPA and at Day 8 influenza post-influenza infection and were downregulated during *A. fumigatus* infection (Figure 5D-E). LGALS9–HAVCR2 (TIM-3) signaling within the T cell compartment showed higher interaction strength during IAPA compared to singular *A. fumigatus infection* (Figure 5F), further supporting this exhaustion signature, as Galectin-9–Tim-3 engagement promotes apoptosis of effector Th1 cells and suppresses T cell–mediated immunity^34^.The TGF-β1–TGFBR1/TGFBR2 interaction, which is pivotal for the induction of FOXP3 expression and differentiation of regulatory T cells^35^, was enriched in the *A. fumigatus* model, consistent with reports that *A. fumigatus* exploits Treg expansion as an immune evasion mechanism^36^ (Figure 5F). Together, these data reveal broad transcriptional dysregulation across T cell, B cell, and NK cell populations during IAPA, including signatures of T cell exhaustion and suppressed Th17 immunity.

**Figure 5.**
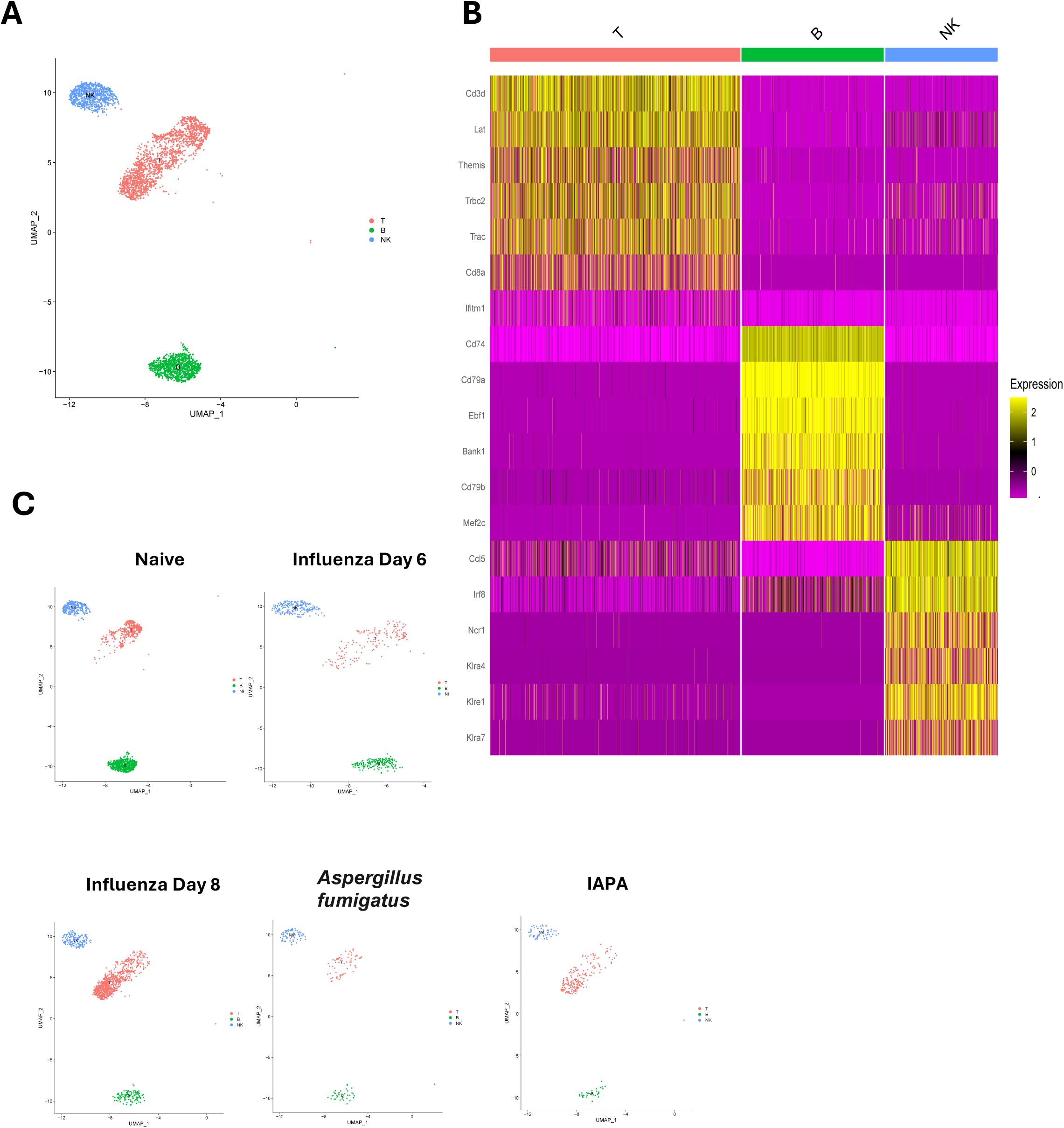
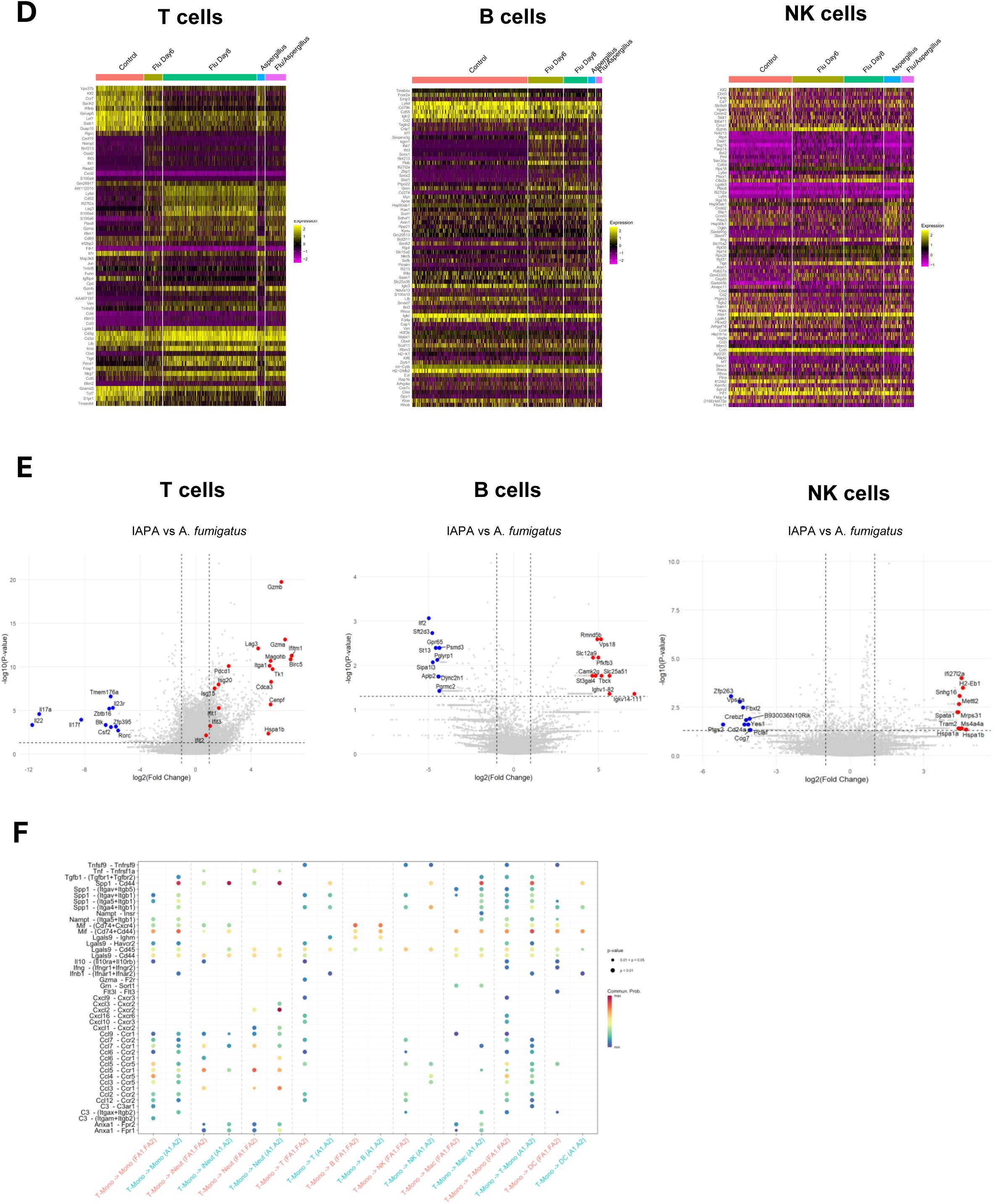
Prior influenza infection disrupts lymphoid antifungal immunity through T cell exhaustion. Influenza Day 6 mice were harvested at day 6; all remaining groups were harvested at day 8. **(A)** UMAP plot of all integrated samples **(B)** Heatmap of biologically relevant differentially expressed genes in each lymphoid cluster **(C)** UMAP plot of all samples, divided by treatment groups **(D)** Differential gene expression in lymphoid clusters (unbiased approach - 10 most upregulated genes and 10 most downregulated genes for each treatment group) **(E)** Volcano plot of the top differentially expressed genes in lymphoid cell populations (Comparing IAPA to *Aspergillus fumigatus* alone) **(F)** Ligand–receptor interactions originating from T Lymphocytes. Dot plot showing inferred ligand–receptor interactions with T lymphocytes as the source cell population, comparing *Aspergillus fumigatus* infection (A1.A2) and IAPA (FA1.FA2) across all target cell types. Dot size reflects p-value significance, and color reflects communication probability.

### Murine IAPA transcriptome recapitulates conserved immune dysfunction signatures identified in human IAPA patients

To assess whether the broad immune remodeling we identified across myeloid, lymphoid, and neutrophil populations reflects pathogenic mechanisms relevant to human disease, we compared our murine transcriptomic data to pathways reported as downregulated in human IAPA bronchoalveolar lavage samples by Feys et al. ^8^ (Figure 6). This cross-species analysis revealed concordance across all eight immune cell types profiled in our murine model, providing strong translational validation of our findings despite differences in species, tissue sampling, and transcriptomic platform.

**Figure 6.**
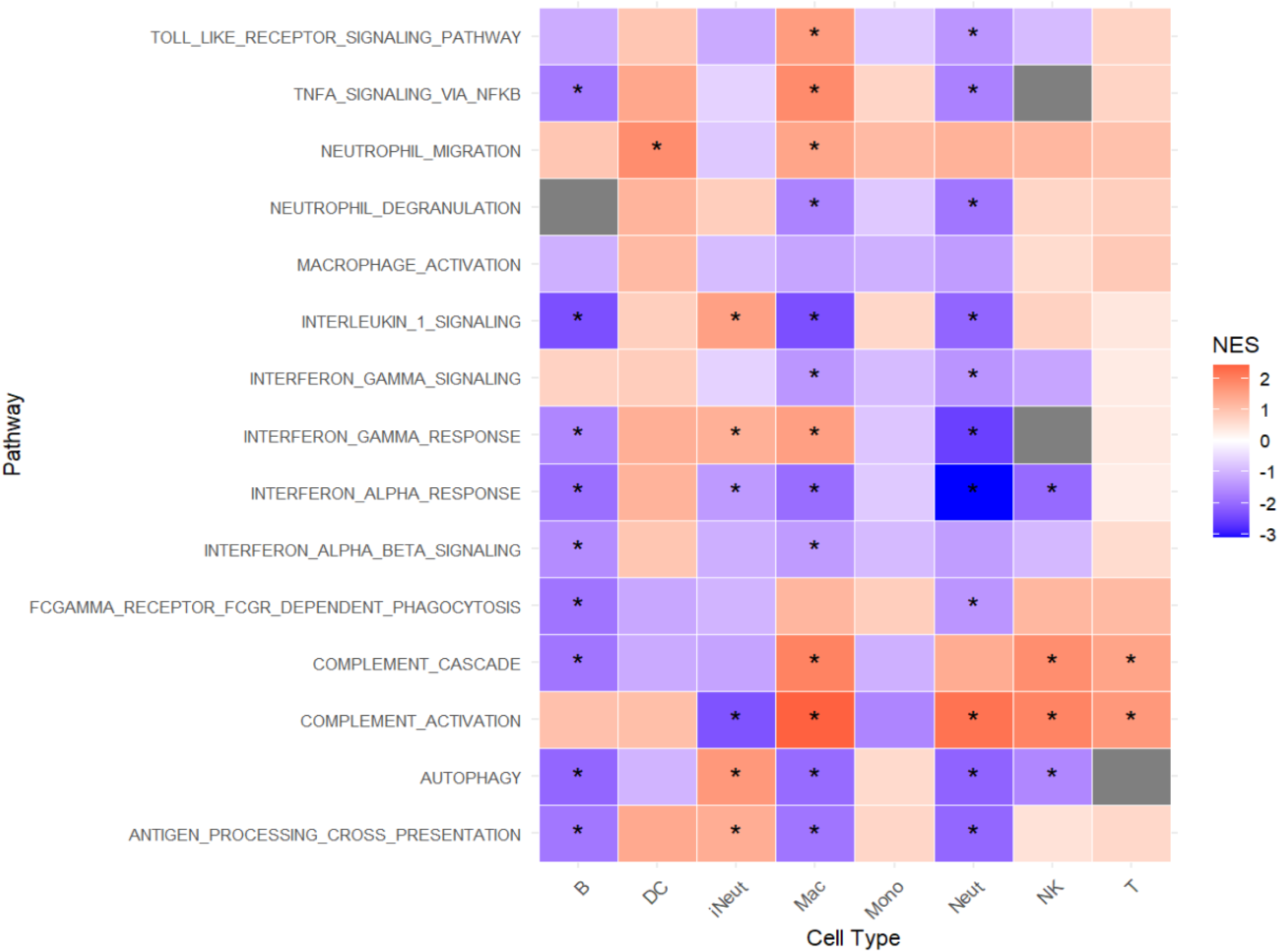
Murine IAPA transcriptome recapitulates conserved immune dysfunction signatures identified in human IAPA patients across all major pulmonary immune cell populations. Gene set enrichment analysis (GSEA) was performed on cell-type-specific differential gene expression comparing IAPA (IAV/*A. fumigatus*) versus singular influenza infection, mirroring the primary biological contrast reported in Feys et al. across all eight major pulmonary immune cell populations identified by scRNA-seq: B cells (B), dendritic cells (DC), inflammatory neutrophils (iNeut), macrophages (Mac), monocytes (Mono), conventional neutrophils (Neut), natural killer cells (NK), and T cells (T). Pathway gene sets were curated to include pathways reported as downregulated in human IAPA bronchoalveolar lavage samples by Feys et al., at either the pathway enrichment or gene level of evidence. Accordingly, a negative NES in our murine dataset reflects directional concordance with the human disease signature. Heatmap color represents the normalized enrichment score (NES) for each pathway-cell type combination: purple-blue/negative NES indicates suppression of the pathway in IAPA relative to influenza infection; orange-red/positive NES indicates relative enrichment of the pathway in IAPA versus influenza infection. Color intensity reflects the magnitude of enrichment or suppression. Gray cells indicate insufficient gene coverage for that pathway within the respective cell type, precluding GSEA computation. Asterisks (*) denote pathways reaching statistical significance at a Benjamini-Hochberg-adjusted p < 0.25, the recommended threshold for GSEA.

These data demonstrate that our murine IAPA model recapitulates the conserved transcriptional defects of human IAPA across myeloid, lymphoid, and neutrophil compartments, and that the immune dysfunction we describe here reflects a biologically conserved response to secondary fungal infection in the post-viral lung.

## Discussion

Dual infection with influenza followed by *A. fumigatus* (IAPA) demonstrated broad remodeling of the pulmonary immune landscape, altering the composition of myeloid, lymphoid, and neutrophil populations relative to single-infection models.

Within the myeloid cell populations, the transcriptional profile of monocytes during IAPA reveals a coordinated suppression of fungal pattern recognition and downstream inflammatory signaling. Downregulation of *Clec4n* (Dectin-2), a principal receptor for *A. fumigatus* galactomannan, suggests impaired upstream fungal recognition in the context of prior influenza infection ^18^. This is compounded by downregulation of *Ptx3*, a soluble pattern recognition receptor that promotes complement activation and opsonization of *A. fumigatus* conidia for phagocytosis by neutrophils and macrophages ^37^, suggesting that both cell-associated and soluble recognition mechanisms may be disrupted in monocytes during IAPA. The clinical significance of PTX3 deficiency in aspergillosis susceptibility is underscored by studies demonstrating that *Ptx3* genetic polymorphisms are independently associated with increased risk of invasive aspergillosis in hematopoietic stem cell transplant recipients, solid organ transplant recipients, and patients with acute leukemia undergoing intensive chemotherapy, with reduced PTX3 expression in neutrophils leading to impaired phagocytosis and fungal clearance ^38–40^. Downstream of pattern recognition, the concurrent downregulation of *Cxcl1* and *Il1b* in monocytes may be linked to impaired neutrophil recruitment and inflammasome activation as functional consequences of this suppressed state. CXCL1 is a critical chemoattractant for neutrophils signaling through CXCR2, and while its reduction in IAPA is consistent with previously published data, the present study identifies monocytes as the specific cellular source of this chemokine suppression^23^. The upregulation of *Ccl8*/MCP-2 in IAPA monocytes represents an intriguing counterpoint to this suppressive pattern; whether this reflects a compensatory recruitment signal or a distinct monocyte activation state during IAPA remains to be determined.

The impact of preceding influenza on macrophage antifungal function has been described in prior publications. Seldeslachts et al. reported a reduction in alveolar macrophage numbers alongside IFN-γ-driven Th17 dysfunction, impairing both the phagocytic and fungicidal capacity of alveolar macrophages against *A. fumigatus* conidia during IAPA ^31^. Consistent with this, our data demonstrate a progressive reduction in macrophage abundance across infection conditions, with the greatest depletion observed during IAPA, suggesting that influenza-driven immune dysregulation contributes to both quantitative and qualitative macrophage defects during coinfection. In contrast, Liu et al. and Lee et al. reported intact antifungal activity of alveolar and interstitial macrophages following antecedent influenza, attributing increased mortality primarily to dysregulated inflammation rather than phagocytic failure^25,28^. Notably, the IAPA models used by different groups vary slightly in the timing of both secondary fungal infection during influenza and the timing of harvest post-fungal infection. Our transcriptional data, showing downregulation of *Tfeb*, *Tyk2*, and *Nfkb1* alongside upregulation of *Il10* and *Tnfaip6*, suggest there may be an M2-skewed, functionally suppressed macrophage state during IAPA, which future studies can address. Downregulation of *Tfeb* is notable given that TFEB and TFE3 are activated in lung phagocytes during *A. fumigatus* infection via Dectin-1/CARD9 signaling, promoting lysosomal biogenesis and nuclear translocation of these transcription factors, and their deficiency impairs macrophage killing of *A. fumigatus* in vitro ^12^. The concurrent downregulation of *Tyk2*, which normally limits IL-10–mediated immunosuppression by restraining the PGE₂–PKA–CREB pathway ^13^, alongside upregulation of *Il10* and *Tnfaip6*, further supports this interpretation.

Interestingly, *Fcgr1* (CD64) and *Fcgr4* (CD16-2), both activating FcγRs that promote antibody-dependent phagocytic and effector responses on myeloid cells ^17,41^ are both highly upregulated antifungal genes in the myeloid cluster, which were upregulated during IAPA compared with *A. fumigatus* infection, in contrast to the broader suppressive pattern. This could represent a compensatory attempt to maintain phagocytic capacity through antibody-dependent mechanisms when innate pattern recognition and inflammatory signaling pathways are compromised.

Within the neutrophil cell populations, a key finding of our analysis is the identification of two transcriptionally distinct populations, conventional and inflammatory neutrophils, with divergent gene expression profiles and effector functions. The inflammatory neutrophil population, characterized by high expression of *Tnf*, *Il1a*, *Cd63*, and *Ccl3*, represents a transcriptionally activated neutrophil state that emerges during infection. The temporal shift from conventional neutrophil-dominant to inflammatory neutrophil-dominant composition between day 6 and day 8 post influenza infection is consistent with progressive neutrophil transcriptional reprogramming during viral infection, reflecting an evolving inflammatory environment when the secondary fungal challenge occurs in our model. Although both conventional and inflammatory neutrophil numbers are decreased during IAPA compared to *A. fumigatus* alone, inflammatory neutrophils are only slightly decreased, while there is a larger, significant reduction in conventional neutrophil numbers during IAPA compared to *A. fumigatus* alone, suggesting influenza-mediated suppression of conventional neutrophil recruitment to the lungs^23,24^. The convergent downregulation of antifungal genes *Nfkb1*, *Syk*, and *Nlrp3* in IAPA neutrophils suggests coordinated suppression of key fungal pattern-recognition signaling nodes. Syk phosphorylation downstream of Dectin-2 drives NF-κB nuclear translocation, proinflammatory cytokine production, and macrophage oxidative burst against *A. fumigatus*^14^. NLRP3 is activated specifically by the hyphal form of *A. fumigatus* and drives inflammasome assembly, ROS production, and caspase-1–mediated IL-1β maturation essential for neutrophil recruitment and antifungal defense^14^. These findings suggest that prior influenza infection may impose a broad transcriptional block on the NF-κB/Syk/NLRP3 axis in neutrophils, constraining their capacity to mount an effective antifungal response against *A. fumigatus*.

The divergent *Lcn2* expression pattern between conventional and inflammatory neutrophils during IAPA has important implications for iron-mediated host defense. Grau et al. recently demonstrated that IAV infection drives a ∼3-fold elevation in pulmonary BALF iron concentrations by day 6 post-infection, accompanied by coordinated induction of iron-sequestration mediators, including LCN2, hepcidin, and ceruloplasmin^42^. Myeloid cells in IAPA lungs showed markedly reduced expression of iron uptake, import, storage, and sequestration pathways, including *Fth1*, *Ltf*, and *Lcn2*, despite this iron-enriched microenvironment, representing a fundamental disconnect between extracellular iron burden and cellular iron-handling capacity^42^. Our data extend this observation to the neutrophil compartment and demonstrate that while inflammatory neutrophils maintain *Lcn2* upregulation consistent with an intact iron-sequestration response, conventional neutrophils show specific *Lcn2* downregulation, suggesting that this iron-regulatory failure is disproportionately concentrated within the conventional neutrophil subset. Notably, the simultaneous upregulation of *Lcn2* and downregulation of *Fth1* in inflammatory neutrophils suggests a dissociation between extracellular iron sequestration and intracellular iron storage within this population, while inflammatory neutrophils may attempt to restrict iron availability to *A. fumigatus* through Lcn2-mediated siderophore binding, their reduced Fth1 expression may leave them unable to safely store excess intracellular iron, potentially rendering them susceptible to iron-mediated oxidative stress and functional exhaustion in the iron-overloaded post-influenza lung. Given that elevated airway iron directly accelerates *A. fumigatus* germination across multiple strains and impairs macrophage antifungal killing^42^, the failure of inflammatory neutrophils to upregulate *Fth1* in the iron-overloaded post-IAV lung may represent a particularly permissive condition for fungal outgrowth, one in which the most activated neutrophil population is simultaneously the most compromised in its ability to limit iron availability to the pathogen. This conventional neutrophil-specific iron-regulatory failure is consistent with the broader pattern of functional impairment within the conventional neutrophil population that we describe throughout this paper. Whether the iron-enriched microenvironment directly drives conventional neutrophil-specific *Lcn2* suppression, or whether this reflects a more general transcriptional suppression program within this population, remains an important question for future investigation.

As a validation of our transcriptomic findings, we observed decreased conventional neutrophil recruitment to the lung in response to *A. fumigatus* challenge in the setting of preceding influenza, coupled with reduced phagocytic uptake of conidia by both conventional and inflammatory neutrophils and reduced killing capacity of conidia by inflammatory neutrophils. The newly identified functional deficits of neutrophil subsets provide important mechanistic data to the IAPA literature. These findings are consistent with other reports of defects in neutrophil killing and phagocytosis ^25,28,29^, supporting the interpretation that neutrophil effector dysfunction is a genuine contributor to IAPA pathogenesis rather than solely reliant on dysregulated inflammation.

Within the lymphoid cell populations, the suppression of Th17 immunity during IAPA, with downregulation of *Il17a*, *Il17f*, *Il22*, *Il23r*, is consistent with prior work identifying decreased Type 17 immune responses during IAPA ^5,31,43^. Our group recently demonstrated that restoration of type 17 immune signaling in our IAPA model is insufficient to provide protection against the increased fungal burden observed ^43^, suggesting that while Type 17 immune suppression is a prominent feature of immune dysregulation during IAPA, it represents one component of a broader immune failure rather than the sole determinant of susceptibility.

The upregulation of *Pdcd1* (PD-1) and *Lag3*, both key regulators of T cell exhaustion and Treg suppressive function ^32,33^, at day 8 post-influenza infection and during IAPA, is consistent with influenza-driven T cell exhaustion persisting into the fungal challenge window. While we did not directly assess T cell functional exhaustion, the transcriptional pattern suggests that prior influenza infection may leave T cells in a partially exhausted state at the time of *A. fumigatus* exposure, potentially limiting the adaptive immune contribution to antifungal defense.

The robust upregulation of interferon-stimulated genes, *Isg15*, *Isg20*, and the IFIT family, in T cells, conventional neutrophils, and inflammatory neutrophils during IAPA compared to *A. fumigatus* infection reflects the persistent antiviral interferon response driven by preceding influenza infection. During influenza infection, ISG15 and ISG20 are strongly induced and play critical roles in limiting viral replication and dissemination^44–46^.Regarding the IFIT family, while these proteins are well-established antiviral effectors^47^, emerging evidence from a systemic Candida model suggests that IFIT2 may play a pathogenic role in antifungal immunity by suppressing neutrophil oxidative responses ^48^. Whether a similar mechanism operates during pulmonary *A. fumigatus* infection in the context of IAPA remains to be determined.

In summary, our findings reveal that preceding influenza infection broadly remodels the pulmonary immune landscape that responds to secondary *A. fumigatus* infection, altering myeloid, lymphoid, and neutrophil populations at the cellular level. Within myeloid cells, we observed coordinated downregulation of fungal pattern recognition receptors, lysosomal biogenesis programs, and key inflammatory signaling. Within lymphoid cells, transcriptional signatures of T cell exhaustion and NK cell dysregulation collectively impair the adaptive immune contribution to antifungal immunity. Within neutrophils, the identification of a transcriptionally and functionally distinct inflammatory neutrophil population, impaired in both fungicidal killing and phagocytic uptake, iron sequestration, and oxidative effector programs, provides a previously unrecognized cellular mechanism for the immune dysregulation observed during IAPA. Taken together, these data support a model in which IAPA susceptibility arises not from a single immunological failure but because of transcriptional changes observed across both innate and adaptive immune cell populations.

## Acknowledgments

We thank Dr. Tobias Hohl (Memorial Sloan Kettering Cancer Center) for generously providing the FLARE conidia used in this study. Work performed in the University of Pittsburgh Single Cell Core Facility (RRID:SCR_025110) and services and instruments used in this project were supported, in part, by the University of Pittsburgh, the Office of the Senior Vice Chancellor for Health Sciences.

## Supplemental Files

**Figure S1: Global cell-cell communication is increased during IAPA**

Total number of inferred ligand–receptor interactions and aggregate interaction strength comparing aspergillus infection and IAPA. IAPA was associated with a greater number of interactions and higher aggregate interaction strength.

**Figure S2: Network topology of intercellular communication during IAPA**

Chord diagrams showing inferred ligand–receptor interaction networks across all immune cell populations during Aspergillus infection and IAPA. Scatter plots showing outgoing versus incoming interaction strength for each cell population during Aspergillus infection and IAPA.

