## Supplemental File 1 for "Broad remodeling of the pulmonary immune landscape occurs during IAPA with specific functional deficits of neutrophil subsets"

#### Table S1

| | Naïve | Flu Day 6 | Flu Day 8 | *Aspergillus fumigatus* | IAPA |
|---|---|---|---|---|---|
| Monocytes | 111 | 613 | 713 | 507 | 908 |
| Macrophages | 525 | 32 | 35 | 50 | 19 |
| Dendritic Cells | 72 | 148 | 77 | 119 | 82 |
| T Cells | 435 | 167 | 855 | 69 | 189 |
| B Cells | 600 | 182 | 123 | 38 | 32 |
| NK Cells | 266 | 215 | 166 | 70 | 54 |
| Neutrophils | 284 | 492 | 283 | 919 | 128 |
| Inflammatory Neutrophils | 1 | 443 | 395 | 1328 | 1025 |

**Table S1 – Immune Cell Subset Abundance and Activation Marker Expression Across Infection Models**

Data represent Absolute cell counts of immune populations obtained from single-cell RNA sequencing of lung immune cells in five experimental conditions: Naïve (PBS control), Influenza Day 6, Influenza Day 8, *Aspergillus fumigatus* infection, and IAPA.

### Figure S1

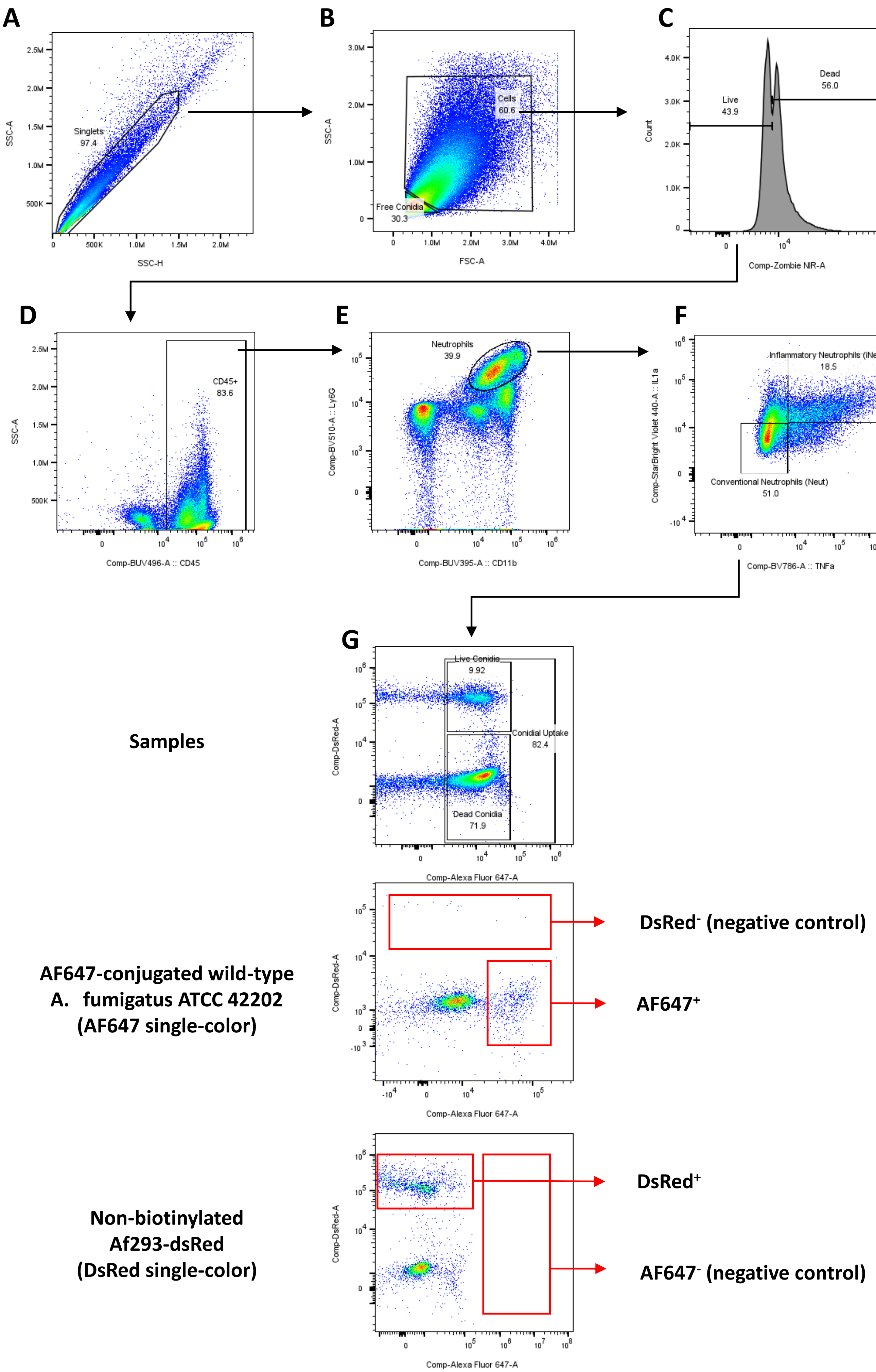
A
Singlets
97.4
SSC-A
SSC-H
B
Cells
60.6
Free Conidia
30.3
SSC-A
FSC-A
C
Live
43.9
Dead
56.0
Count
Comp-Zombie NIR-A
D
CD45+
83.6
SSC-A
Comp-BUV496-A :: CD45
E
Neutrophils
39.9
Comp-BV510-A :: Ly6G
Comp-BUV395-A :: CD11b
F
Inflammatory Neutrophils (iNeut)
18.5
Conventional Neutrophils (Neut)
51.0
Comp-StarBright Violet 440-A :: IL1a
Comp-BV786-A :: TNFa
G
Live Conidia
9.92
Conidial Uptake
82.4
Dead Conidia
71.9
Comp-DsRed-A
Comp-Alexa Fluor 647-A
Samples
AF647-conjugated wild-type
A. fumigatus ATCC 42202
(AF647 single-color)
DsRed- (negative control)
AF647+
Non-biotinylated
Af293-dsRed
(DsRed single-color)
DsRed+
AF647- (negative control)

**Figure S1 – Flow Cytometry Gating Strategy for Neutrophil Identification and Conidial Uptake**

Lung cells were analyzed 48 hours following infection with $1\times10^8$ *A. fumigatus* FLARE conidia. Lung single-cell suspensions were prepared as described in the Methods. **(A)** Singlets were identified by SSC-H vs SSC-A gating. **(B)** Free *Aspergillus fumigatus* conidia were gated as $FSC^{lo}$ $SSC^{lo}$ events and excluded; immune cells were selected from the remaining population. **(C)** Live/dead discrimination was performed using Zombie NIR. **(D)** $CD45^+$ leukocytes were gated. **(E)** Neutrophils were identified as $CD45^+$ $Ly6G^+$ $CD11b^+$ cells. **(F)** Within the neutrophil gate, inflammatory neutrophils (iNeut) were defined by high expression of IL-1α and TNF (IL-$1\alpha^{high}TNF\alpha^{high}$), while conventional neutrophils (Neut) were characterized by low expression of these markers (IL-$1\alpha^{lo}TNF\alpha^{lo}$). **(G)** Conidial uptake was determined by AF647 positivity. $DsRed^+$ $AF647^+$ cells represent neutrophils containing live conidia, whereas $DsRed^-$ $AF647^+$ cells indicate neutrophils containing dead conidia. Each flowcytometry experimental run included two compensation controls: a mouse challenged with non-biotinylated Af293-dsRed conidia to serve as the DsRed single-color reference, and a mouse challenged with AF647-conjugated wild-type A. fumigatus ATCC 42202 to serve as the AF647 single-color reference. These controls were processed in parallel with experimental samples to ensure accurate spectral unmixing and population gating.

### Figure S2

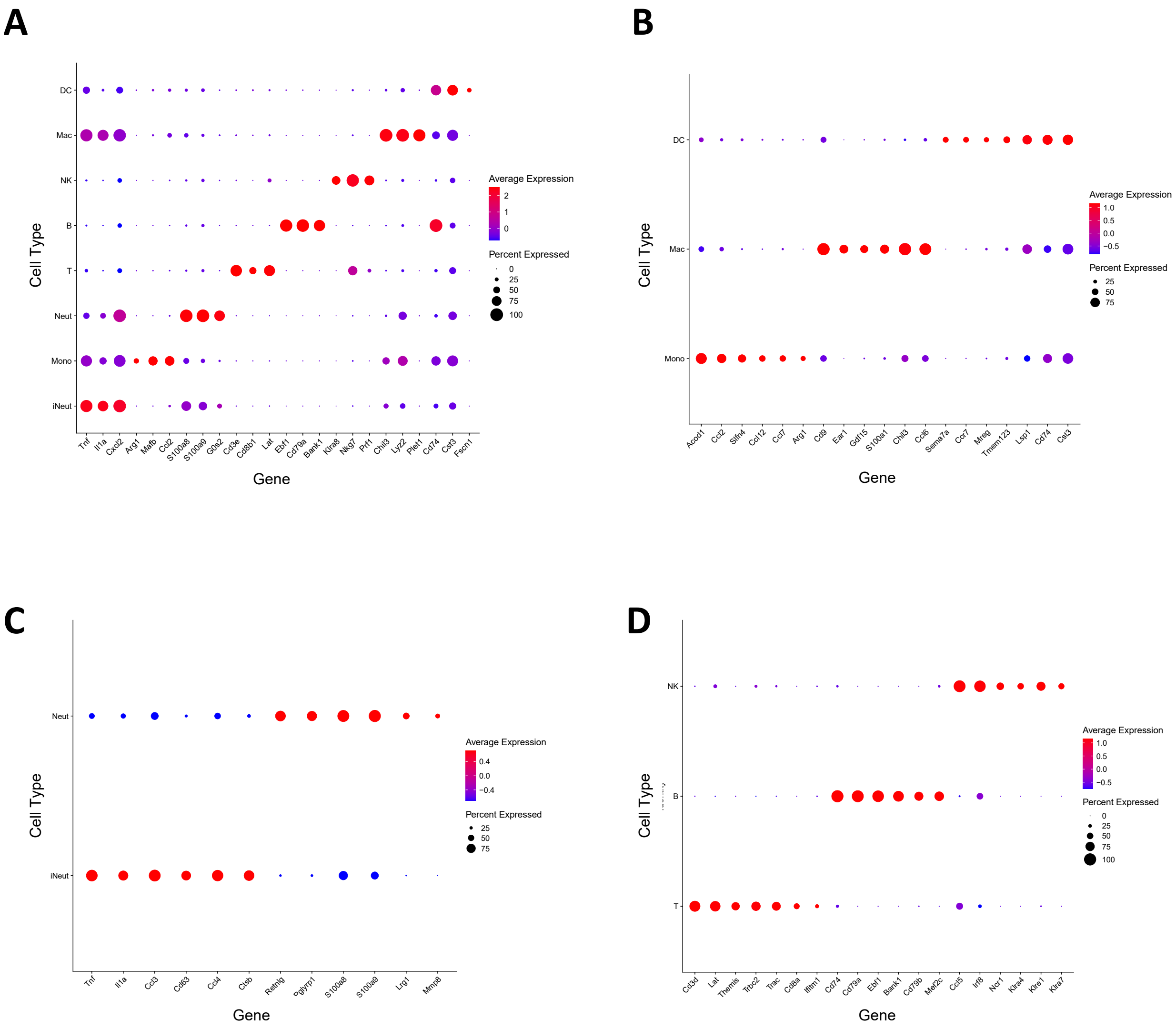
A
Cell Type
DC
Mac
NK
B
T
Neut
Mono
iNeut
Tnf
Il1a
Cxcl2
Arg1
Mafb
Ccl2
S100a8
S100a9
G0s2
Cd3e
Cd8b1
Lat
Ebf1
Cd79a
Bank1
Klra8
Nkg7
Prf1
Chil3
Lyz2
Plet1
Cd74
Cst3
Fscn1
Gene
Average Expression
2
1
0
Percent Expressed
0
25
50
75
100
B
Cell Type
DC
Mac
Mono
Acod1
Ccl2
Slfn4
Ccl12
Ccl7
Arg1
Cd9
Ear1
Gdf15
S100a1
Chil3
Ccl6
Sema7a
Ccr7
Mreg
Tmem123
Lsp1
Cd74
Cst3
Gene
Average Expression
1.0
0.5
0.0
−0.5
Percent Expressed
25
50
75
C
Cell Type
Neut
iNeut
Tnf
Il1a
Ccl3
Cd63
Ccl4
Cstb
Retnlg
Pglyrp1
S100a8
S100a9
Lrg1
Mmp8
Gene
Average Expression
0.4
0.0
−0.4
Percent Expressed
25
50
75
D
Cell Type
NK
B
T
Cd3d
Lat
Themis
Trbc2
Trac
Cd8a
Ifitm1
Cd74
Cd79a
Ebf1
Bank1
Cd79b
Mef2c
Ccl5
Irf8
Ncr1
Klra4
Klre1
Klra7
Gene
Average Expression
1.0
0.5
0.0
−0.5
Percent Expressed
0
25
50
75
100

**Figure S2. Dot plots showing marker gene expression across immune cell populations.**

**(A)** Dot plot showing expression of canonical lineage marker genes across all eight identified immune cell populations.

**(B)** Dot plot showing expression of marker genes within myeloid cell clusters (monocytes, macrophages, and dendritic cells).

**(C)** Dot plot showing expression of marker genes within neutrophil clusters (conventional neutrophils and inflammatory neutrophils).

**(D)** Dot plot showing expression of marker genes within lymphoid cell clusters (T cells, B cells, and NK cells).

Dot size reflects the proportion of cells expressing each gene within the indicated cell type, and color intensity reflects the scaled average expression level.

**Figure S3. Transcriptional landscape of pulmonary immune cell populations during IAPA.**

**(A)** Heatmap of all differentially expressed genes in each myeloid cluster (unbiased approach)

**(B)** Heatmap of all differentially expressed genes in each lymphoid cluster (unbiased approach)

**(C)** Heatmap of all differentially expressed genes in each neutrophil cluster (unbiased approach)

### Figure S4

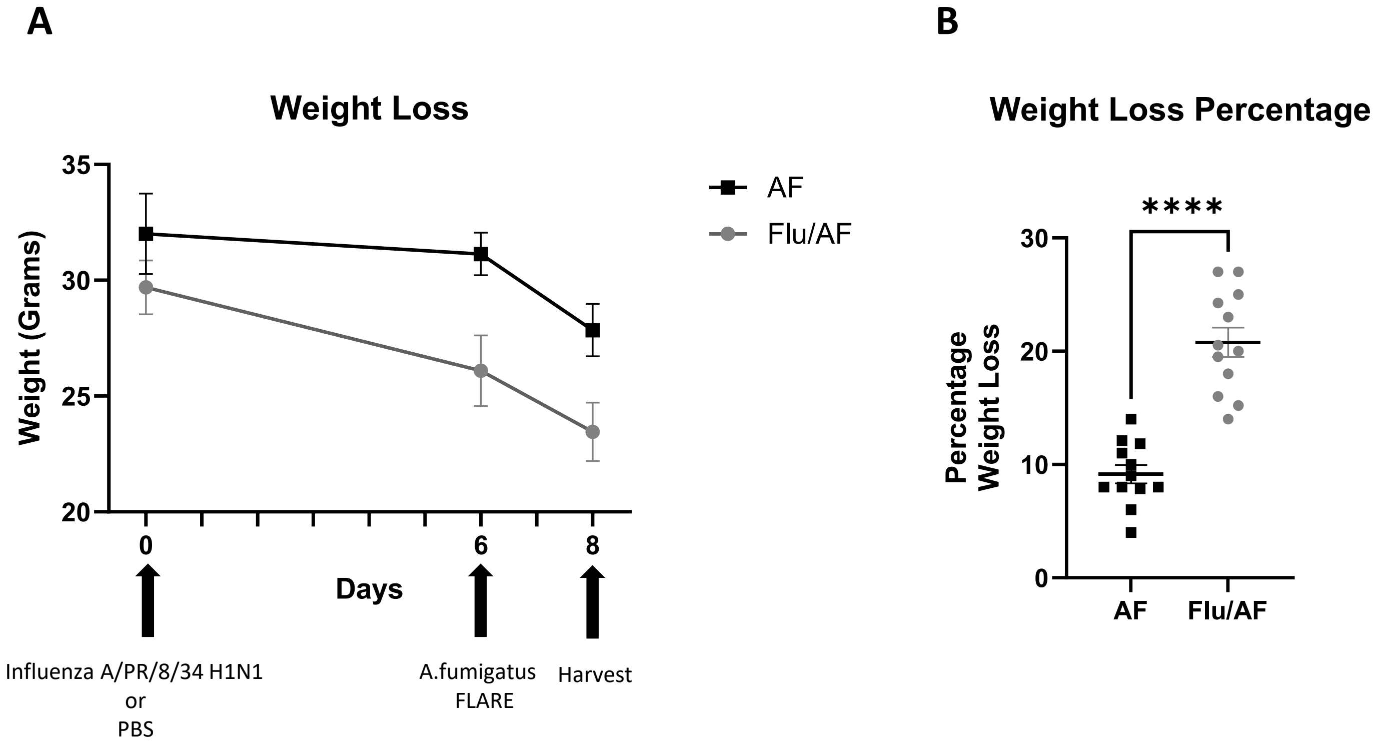
A
Weight Loss
35
30
25
20
Weight (Grams)
AF
Flu/AF
0
6
8
Days
Influenza A/PR/8/34 H1N1
or
PBS
A.fumigatus
FLARE
Harvest
B
Weight Loss Percentage
****
30
20
10
0
Percentage
Weight Loss
AF
Flu/AF

**Figure S4. IAPA is associated with greater weight loss compared to singular *A. fumigatus* infection.**

**(A)** Body weight (grams) of mice at day 0 (prior to influenza A/PR/8/34 H1N1 or PBS inoculation), day 6 (prior to *A. fumigatus* FLARE conidial challenge), and day 8 (harvest) in mice receiving *A. fumigatus* alone (AF) or Flu/AF. Data are presented as mean ± SEM.

**(B)** Percentage weight loss from baseline (day 0) to harvest (day 8) in AF and Flu/AF groups.

Data were compiled from two independent experiments and are presented as mean ± SEM. ****$p<0.0001$ by unpaired t-test.
