## Supplemental File 2 for "Broad remodeling of the pulmonary immune landscape occurs during IAPA with specific functional deficits of neutrophil subsets"

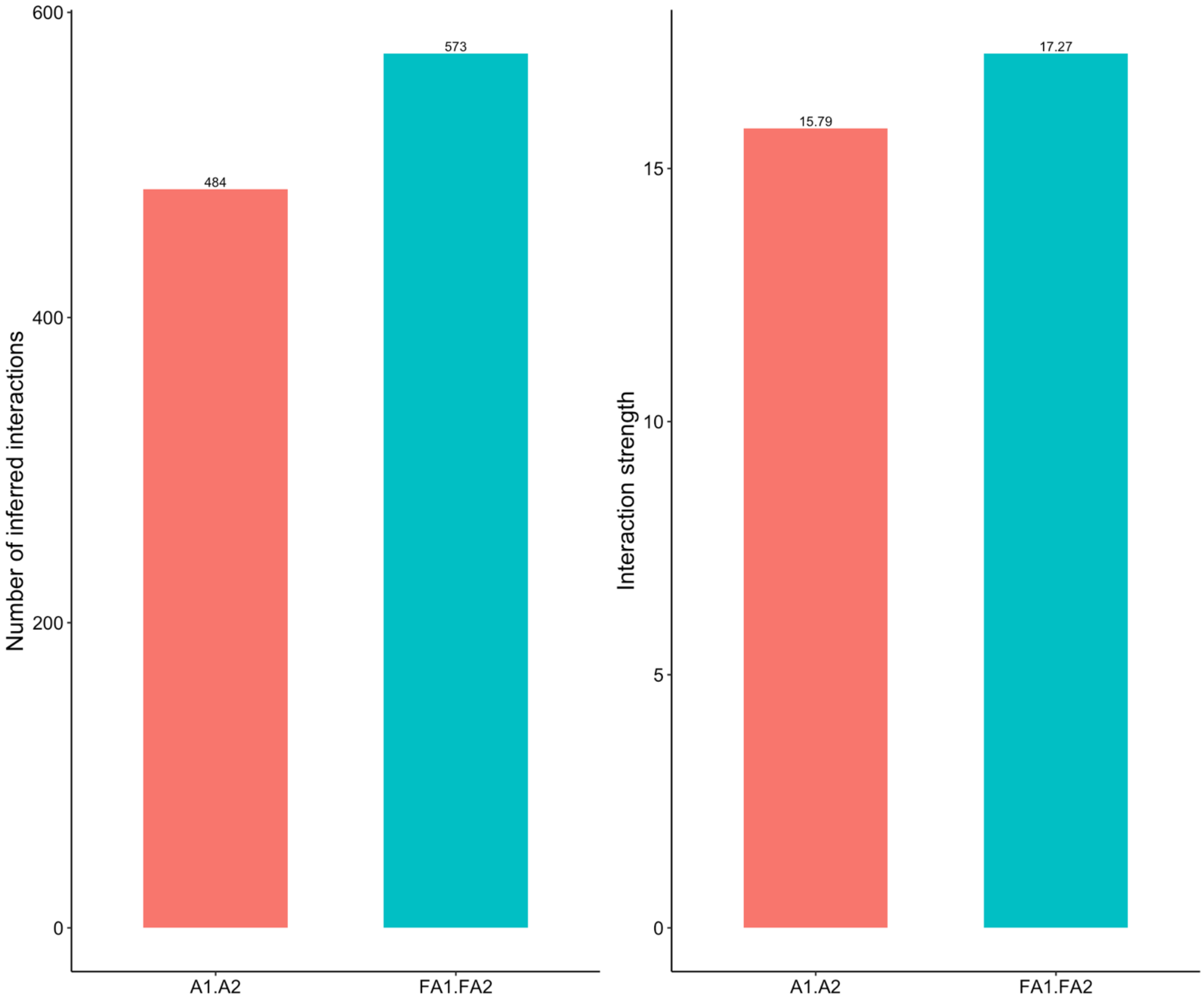


**Figure S1 — Global cell-cell communication is increased during IAPA** Bar charts showing the total number of inferred ligand–receptor interactions (left) and aggregate interaction strength (right) comparing aspergillus infection (A1.A2) and IAPA (FA1.FA2). IAPA was associated with a greater number of interactions (573 vs 484) and higher aggregate interaction strength (17.27 vs 15.79).

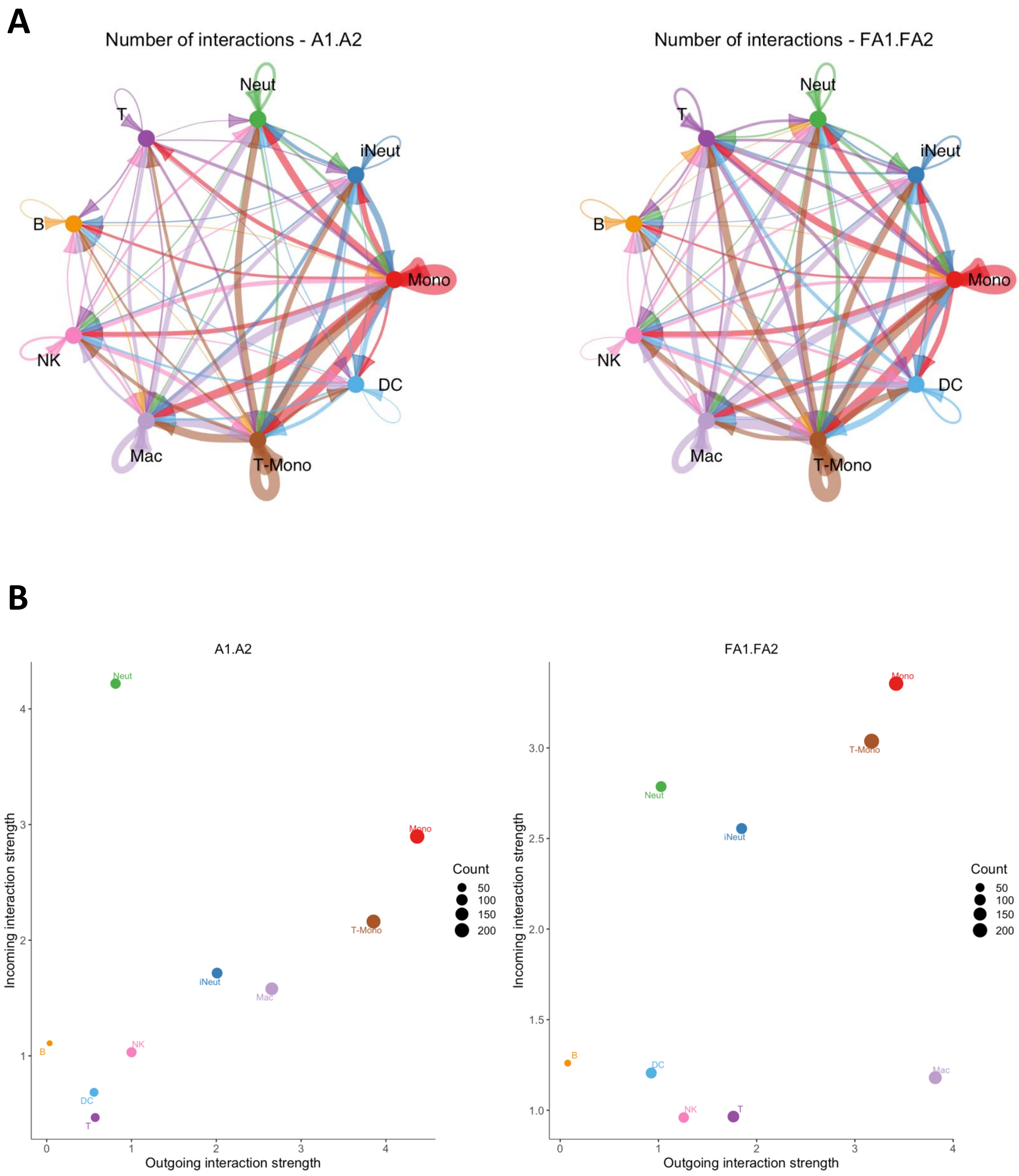

A
Number of interactions - A1.A2
Number of interactions - FA1.FA2
Neut
T
iNeut
B
Mono
NK
DC
Mac
T-Mono
B
A1.A2
FA1.FA2
Incoming interaction strength
Outgoing interaction strength
Count
50
100
150
200

**Figure S2 — Network topology of intercellular communication during IAPA**
**(A)** Chord diagrams showing inferred ligand–receptor interaction networks across all immune cell populations during Aspergillus infection (A1.A2, left) and IAPA (FA1.FA2, right). Arrow thickness indicates interaction strength.
**(B)** Scatter plots showing outgoing versus incoming interaction strength for each cell population during Aspergillus infection (left) and IAPA (right). Dot size reflects the number of interactions.

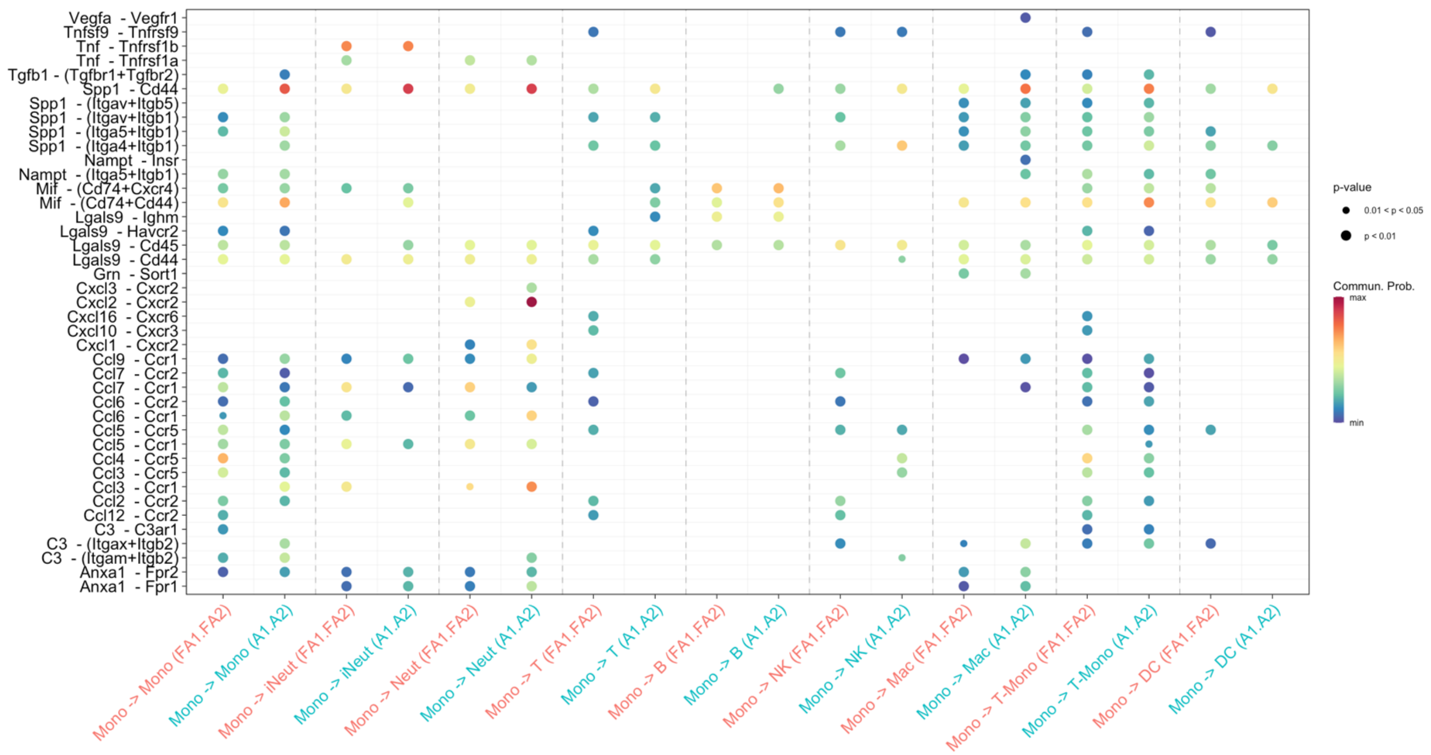


**Figure S3 — Ligand–receptor interactions originating from monocytes.** Dot plot showing inferred ligand–receptor interactions with monocytes as the source cell population, comparing aspergillus infection (A1.A2) and IAPA (FA1.FA2) across all target cell types. Dot size reflects p-value significance, and color reflects communication probability.

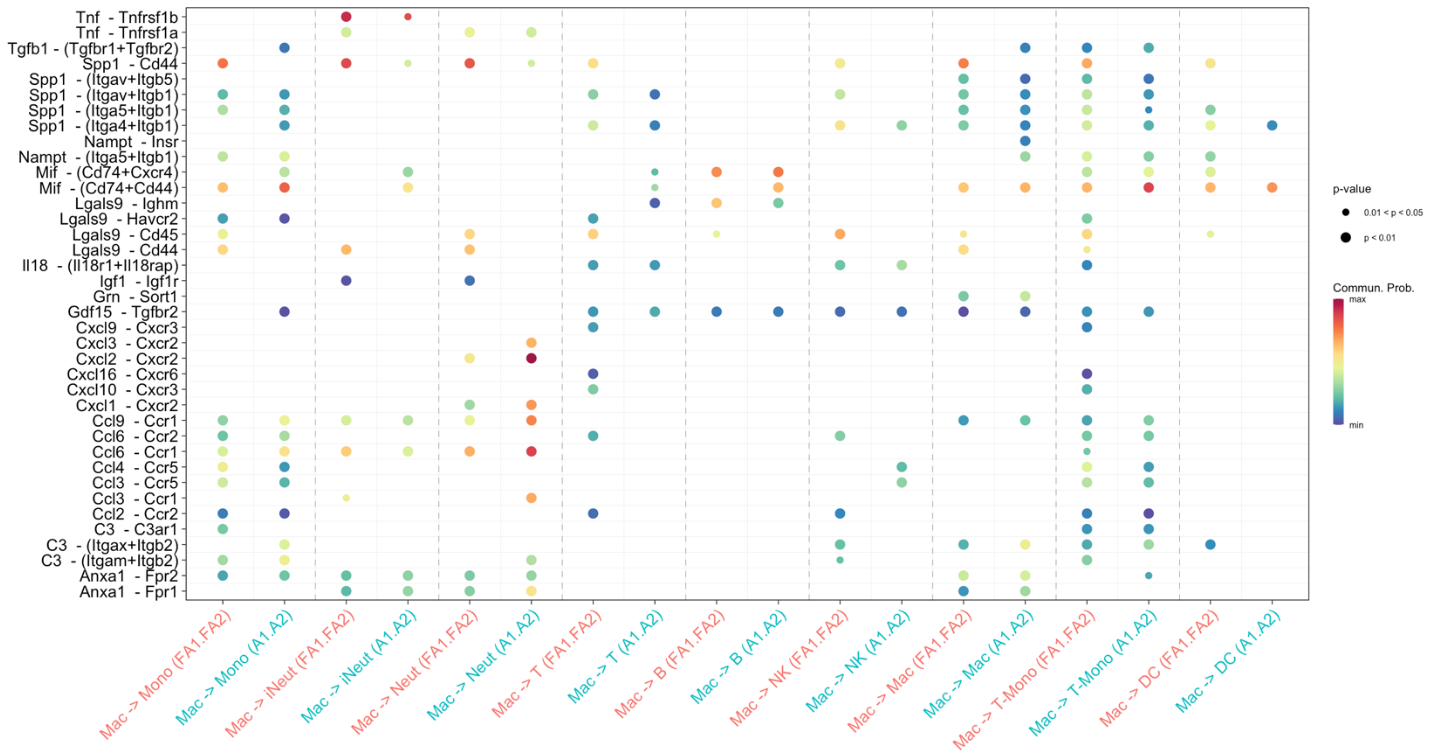


**Figure S4 — Ligand–receptor interactions originating from Macrophages.** Dot plot showing inferred ligand–receptor interactions with macrophages as the source cell population, comparing aspergillus infection (A1.A2) and IAPA (FA1.FA2) across all target cell types. Dot size reflects p-value significance, and color reflects communication probability.

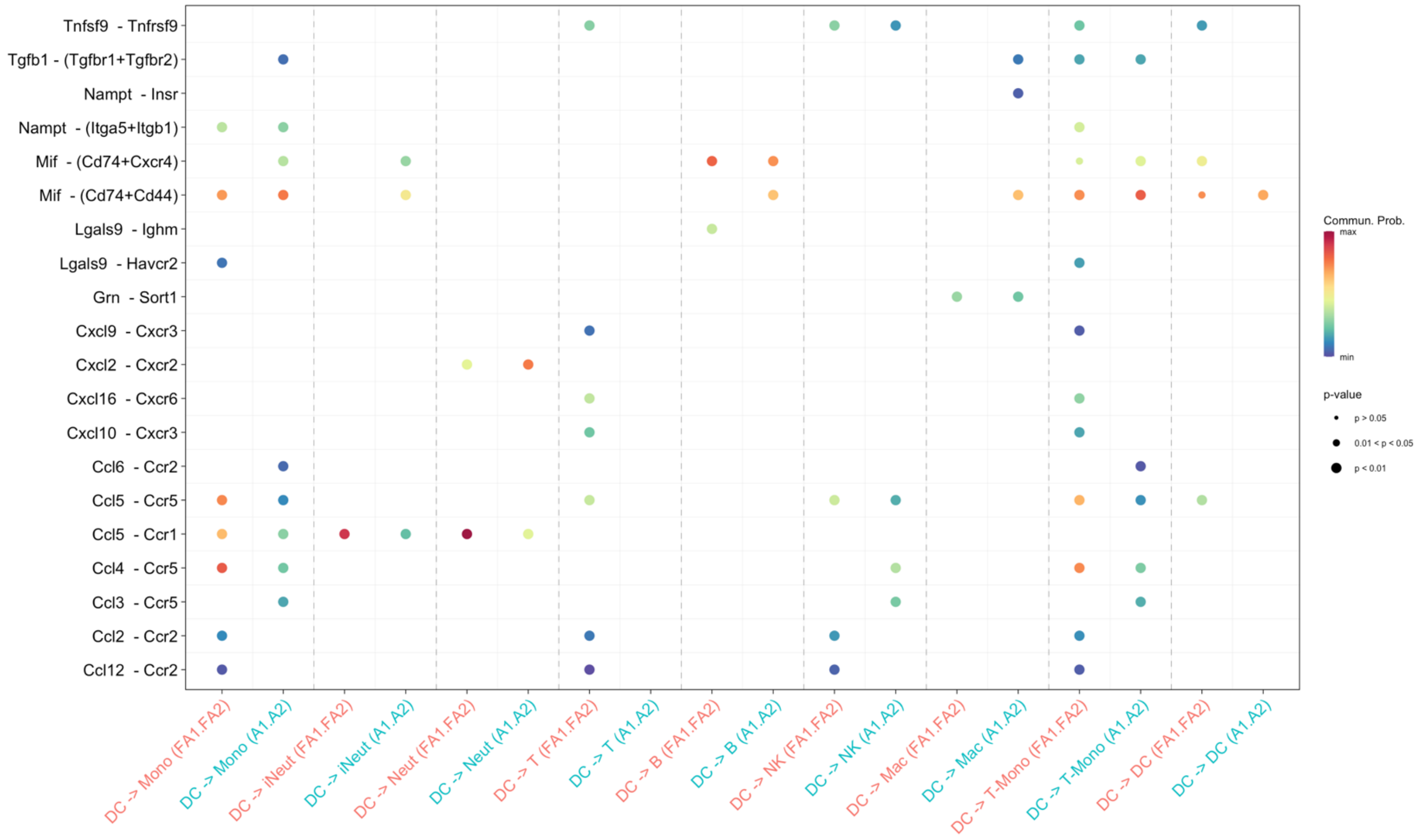


**Figure S5 — Ligand–receptor interactions originating from Dendritic Cells.** Dot plot showing inferred ligand–receptor interactions with dendritic Cells as the source cell population, comparing aspergillus infection (A1.A2) and IAPA (FA1.FA2) across all target cell types. Dot size reflects p-value significance, and color reflects communication probability.

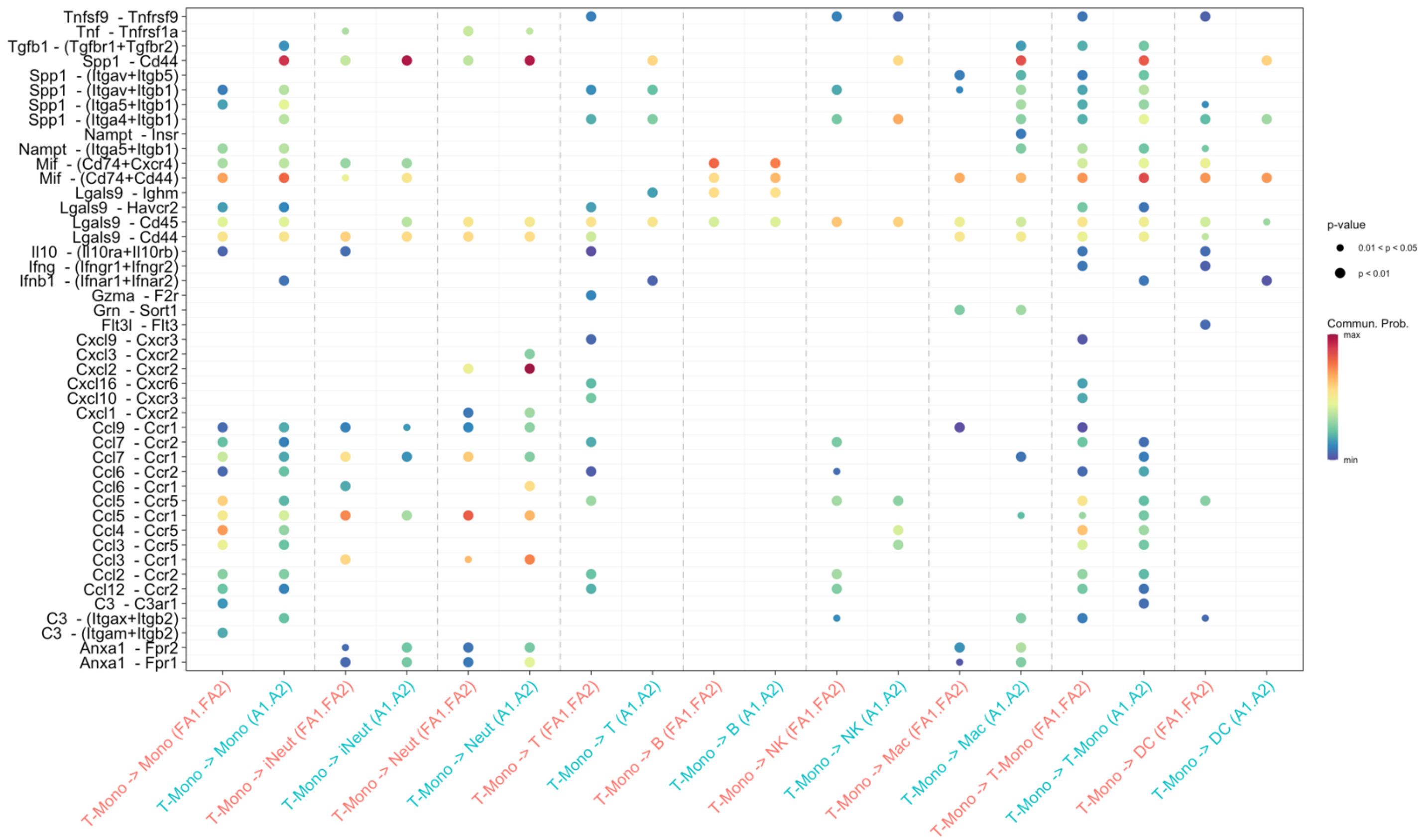


**Figure S6 — Ligand–receptor interactions originating from T Lymphocytes.** Dot plot showing inferred ligand–receptor interactions with T lymphocytes as the source cell population, comparing aspergillus infection (A1.A2) and IAPA (FA1.FA2) across all target cell types. Dot size reflects p-value significance, and color reflects communication probability.

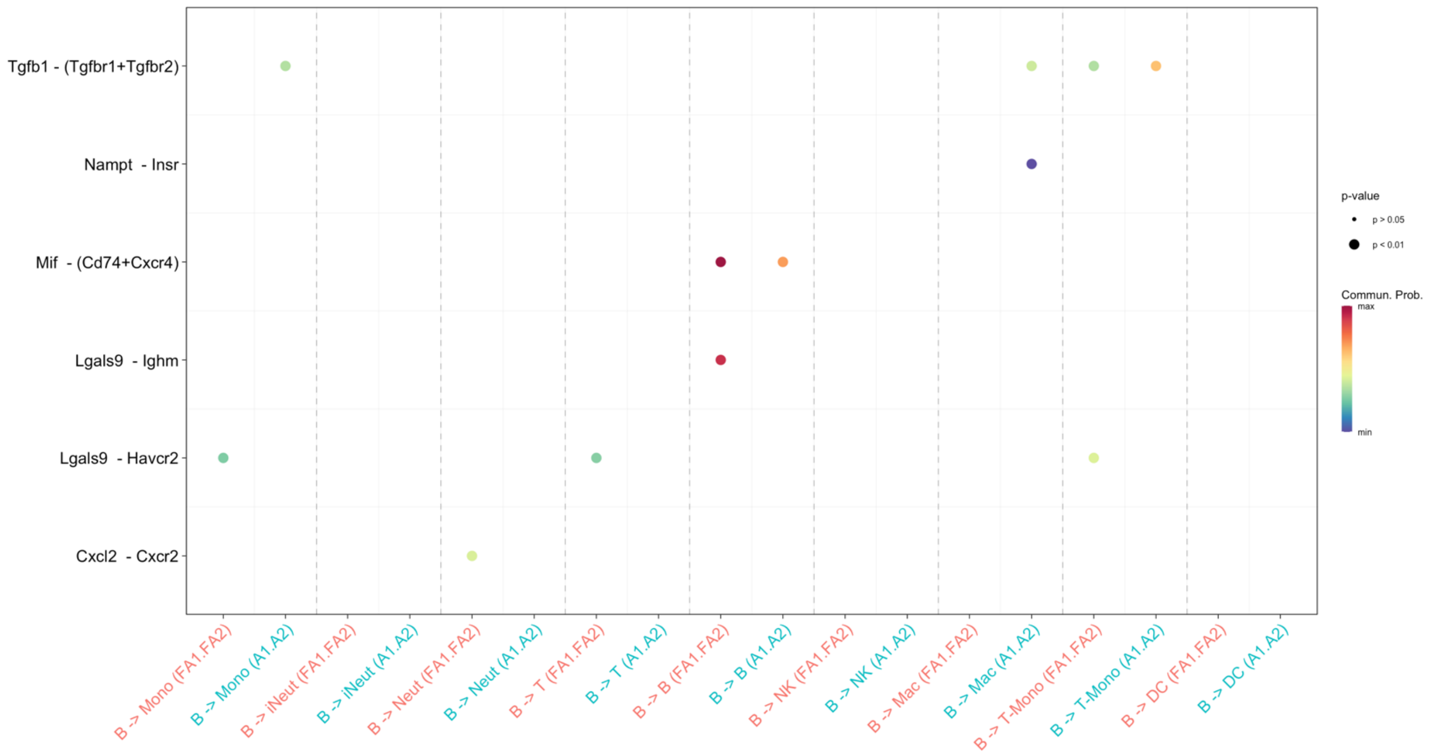


**Figure S7 — Ligand–receptor interactions originating from B Lymphocytes.** Dot plot showing inferred ligand–receptor interactions with B lymphocytes as the source cell population, comparing aspergillus infection (A1.A2) and IAPA (FA1.FA2) across all target cell types. Dot size reflects p-value significance, and color reflects communication probability.

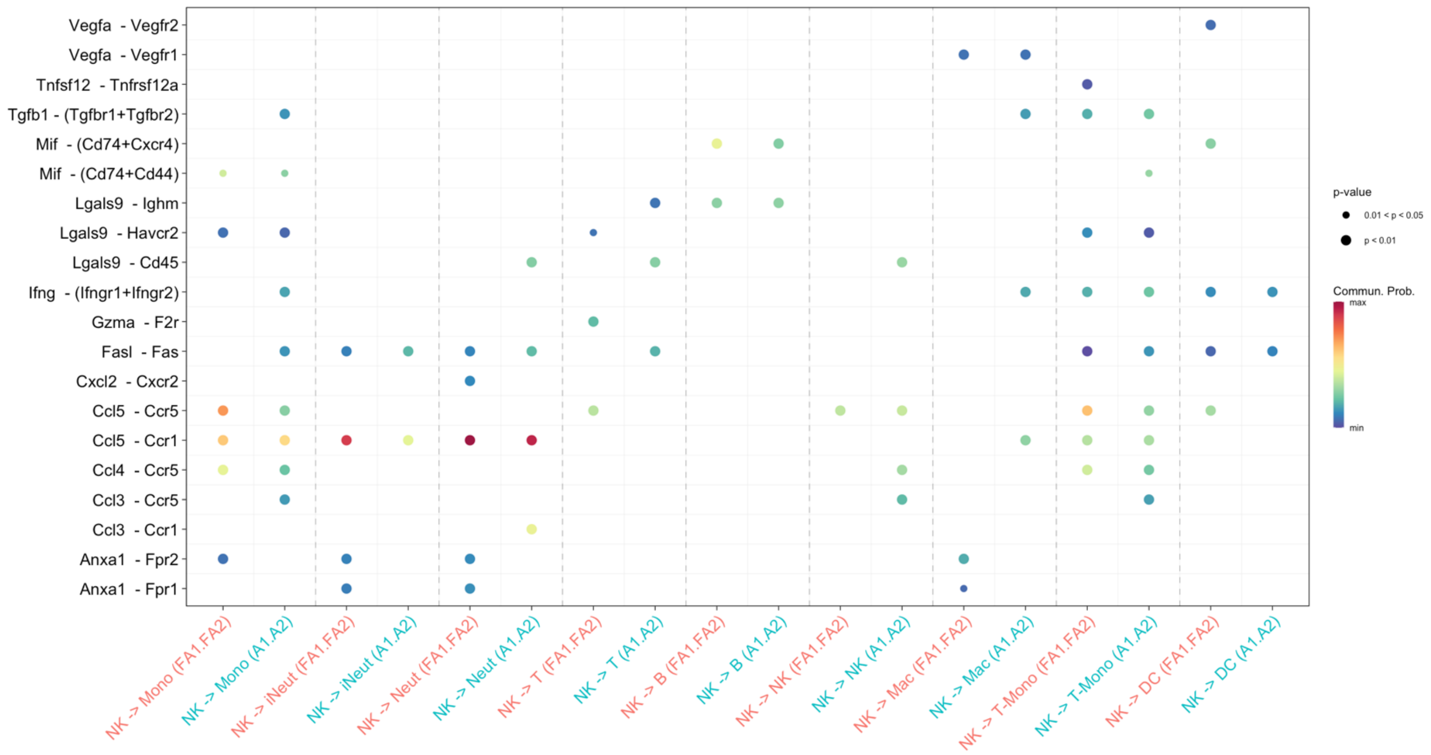


**Figure S8 — Ligand–receptor interactions originating from NK Cells.** Dot plot showing inferred ligand–receptor interactions with NK cells as the source cell population, comparing aspergillus infection (A1.A2) and IAPA (FA1.FA2) across all target cell types. Dot size reflects p-value significance, and color reflects communication probability.

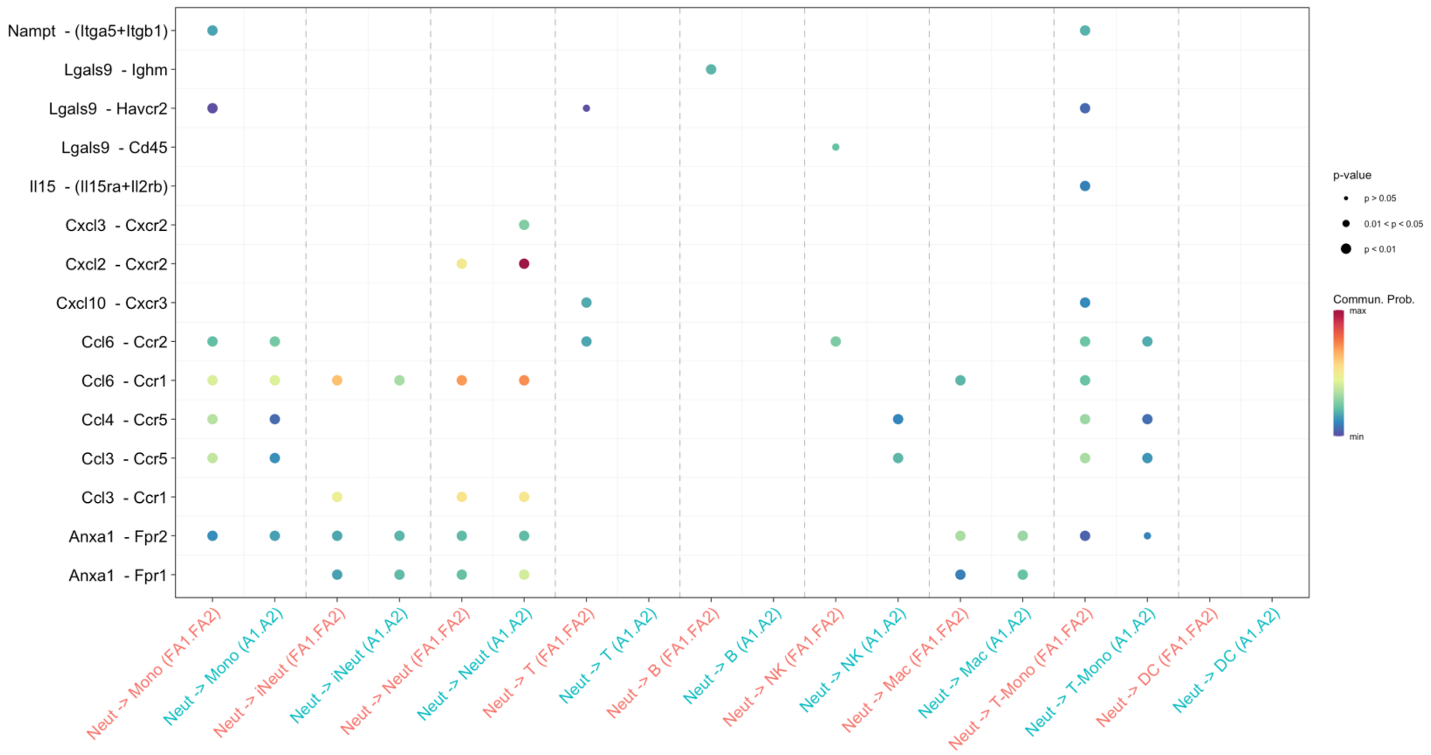


**Figure S9 — Ligand–receptor interactions originating from Conventional Neutrophils (Neut).** Dot plot showing inferred ligand–receptor interactions with Neut as the source cell population, comparing aspergillus infection (A1.A2) and IAPA (FA1.FA2) across all target cell types. Dot size reflects p-value significance, and color reflects communication probability.

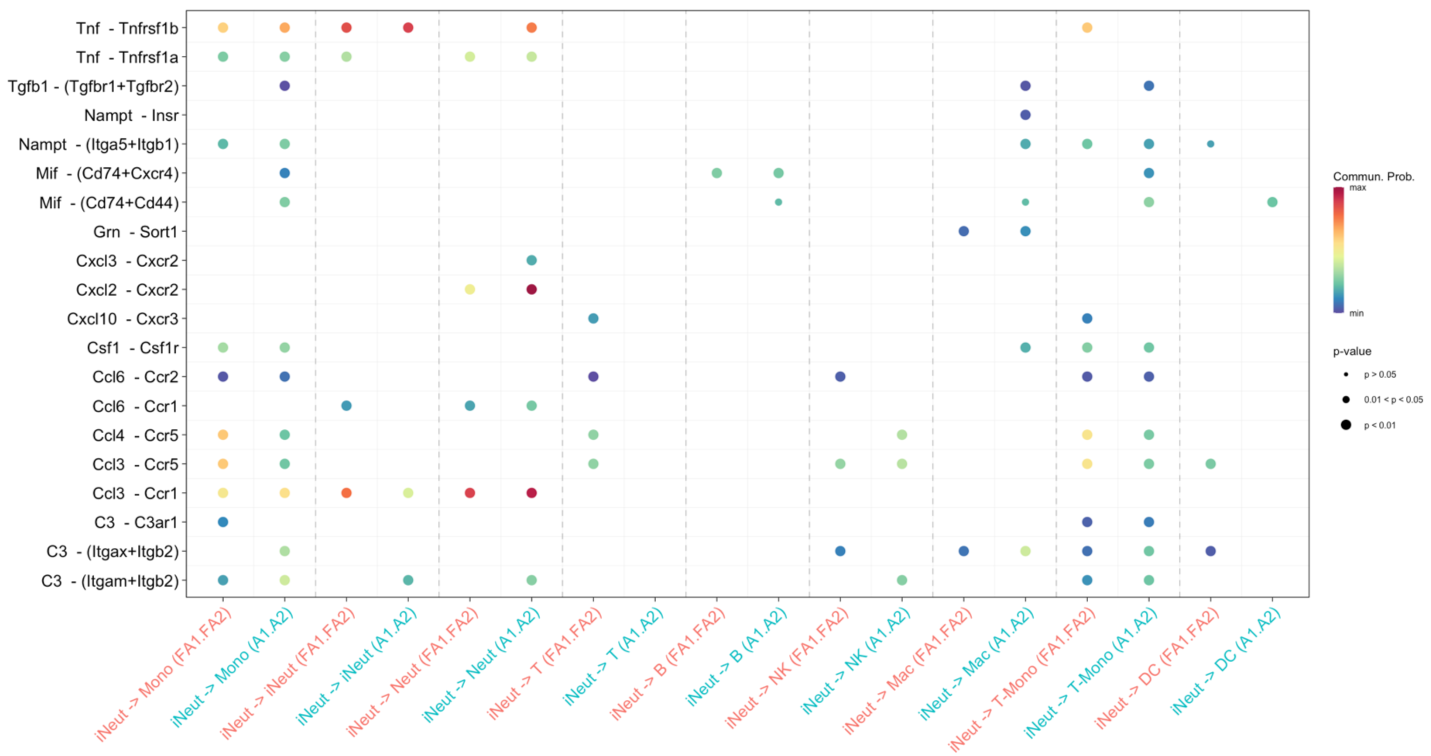


**Figure S10 — Ligand–receptor interactions originating from Inflammatory Neutrophils (iNeut).** Dot plot showing inferred ligand–receptor interactions with iNeut as the source cell population, comparing aspergillus infection (A1.A2) and IAPA (FA1.FA2) across all target cell types. Dot size reflects p-value significance, and color reflects communication probability.
